# Molecular, spatial and projection diversity of neurons in primary motor cortex revealed by in situ single-cell transcriptomics

**DOI:** 10.1101/2020.06.04.105700

**Authors:** Meng Zhang, Stephen W. Eichhorn, Brian Zingg, Zizhen Yao, Hongkui Zeng, Hongwei Dong, Xiaowei Zhuang

**Affiliations:** Howard Hughes Medical Institute, Harvard University, Cambridge, MA 02138, USA; Department of Chemistry and Chemical Biology, Harvard University, Cambridge, MA 02138, USA; Department of Physics, Harvard University, Cambridge, MA 02138, USA; Center for Integrative Connectomics, Mark and Mary Stevens Neuroimaging and Informatics Institute, Keck School of Medicine of USC, University of Southern California, Los Angeles, CA, 90095, USA; Department of Neurology, Keck School of Medicine of USC, University of Southern California, Los Angeles, CA, 90095, USA; Allen Institute for Brain Science, Seattle, WA 98109, USA

**Author notes:** Correspondence should be addressed to (X.Z.).

## Abstract

A mammalian brain is comprised of numerous cell types organized in an intricate manner to form functional neural circuits. Single-cell RNA sequencing provides a powerful approach to identify cell types based on their gene expression profiles and has revealed many distinct cell populations in the brain^1-3^. Single-cell epigenomic profiling^4,5^ further provides information on gene-regulatory signatures of different cell types. Understanding how different cell types contribute to brain function, however, requires knowledge of their spatial organization and connectivity, which is not preserved in sequencing-based methods that involve cell dissociation^3,6^. Here, we used an in situ single-cell transcriptome-imaging method, multiplexed error-robust fluorescence in situ hybridization (MERFISH)^7^, to generate a molecularly defined and spatially resolved cell atlas of the mouse primary motor cortex (MOp). We profiled ∼300,000 cells in the MOp, identified 95 neuronal and non-neuronal cell clusters, and revealed a complex spatial map in which not only excitatory neuronal clusters but also most inhibitory neuronal clusters adopted layered organizations. Notably, intratelencephalic (IT) cells, the largest branch of neurons in the MOp, formed a continuous spectrum of cells with gradual changes in both gene expression profiles and cortical depth positions in a highly correlated manner. Furthermore, we integrated MERFISH with retrograde tracing to probe the projection targets for different MOp neuronal cell types and found that projections of MOp neurons to other cortical regions formed a many-to-many network with each target region receiving input preferentially from a different composition of IT clusters. Overall, our results provide a high-resolution spatial and projection map of molecularly defined cell types in the MOp. We anticipate that the imaging platform described here can be broadly applied to create high-resolution cell atlases of a wide range of systems.

## Main

The cerebral cortex is a highly-organized structure comprised of distinct regions that support different sensory, motor, and cognitive functions. Known for its distinctive laminar structure, the cortex is delineated into six layers (L1-L6) based on cytoarchitectural features. Various cortical regions are interconnected with each other and with other brain region to form functional neural circuits^8-10^. Classification of neuronal cell types is central to deciphering the complexity of these circuits^2,11-13^. The glutamatergic excitatory neurons in the cortex are often classified by their projection properties into, for example, intratelencephalic (IT) neurons, sub-cerebral projection neurons (or pyramidal tract neurons), and cortico-thalamic (CT) projection neurons^14,15^. The GABAergic inhibitory neurons can be divided based on their developmental origin into caudal ganglionic eminence (CGE) derived and medial ganglionic eminence (MGE) derived neurons, and can also be classified by prominent marker genes such as Parvalbumin (Pvalb), Somatostatin (Sst), and Vasoactive intestinal polypeptide (Vip)^16-18^. Recent single-cell transcriptomics studies have revealed an extraordinarily high diversity of cells in the brain^19-21^ and reported dozens to a hundred cell types within individual cortical regions^22-24^. However, a high-resolution map of the spatial organization and connectivity of different cell types in the cortex, which is essential to understanding cortical circuits, is still missing.

Recently, a number of spatially resolved transcriptomics methods have been developed, including both in situ imaging-based transcriptomics methods with single-cell resolution^7,25-29^ and methods based on spatially resolved RNA capture followed by sequencing^30,31^. Among these, multiplexed error-robust fluorescence in situ hybridization (MERFISH) is a single-cell transcriptome imaging method, which massively multiplexes single-molecule FISH^32,33^ using error-robust barcoding, combinatorial labeling and sequential imaging^7^. MERFISH allows simultaneous imaging of hundreds to thousands of genes in individual cells with high detection efficiency both in cultured cells^7,34^ and in brain tissue slices^35,36^. Here we used MERFISH to perform in situ gene expression profiling of individual cells, identifying distinct cell populations and mapping their spatial organization in the mouse primary motor cortex (MOp), a region designated by the BRAIN Initiative Cell Census Network (BICCN)^37^ as the initial target for comprehensive cell mapping. Furthermore, we developed an approach to integrate MERFISH with retrograde tracing, and used this approach to determine the compositions and spatial distributions of MOp neurons that project to several cortical regions.

### Single-cell gene expression profiling and cell type identification of the mouse MOp by MERFISH

We selected a panel of 258 genes for MERFISH imaging (Figure 1a) by combining three distinct approaches: (i) 62 canonical marker genes for major neuronal and non-neuronal cell types in the cortex were selected based on prior knowledge; (ii) 168 genes were selected based on pair-wise differential gene expression analysis on the neuronal clusters identified by a concurrent single-cell / single-nucleus RNA-sequencing (sc/snRNA-seq) study reported in a companion BICCN paper^38^; (iii) a set of genes (50 for glutamatergic and 50 for GABAergic neuronal clusters) were selected which contained the highest mutual information^39,40^ among the clusters identified by sc/snRNA-seq. The overlapping gene lists generated by these three approaches were combined to form a panel of 258 genes total, in which 242 were imaged using MERFISH, and the remaining 16 genes, which were either relatively short or highly expressed, were measured in eight sequential rounds of a two-color FISH following MERFISH.

**Figure 1.**
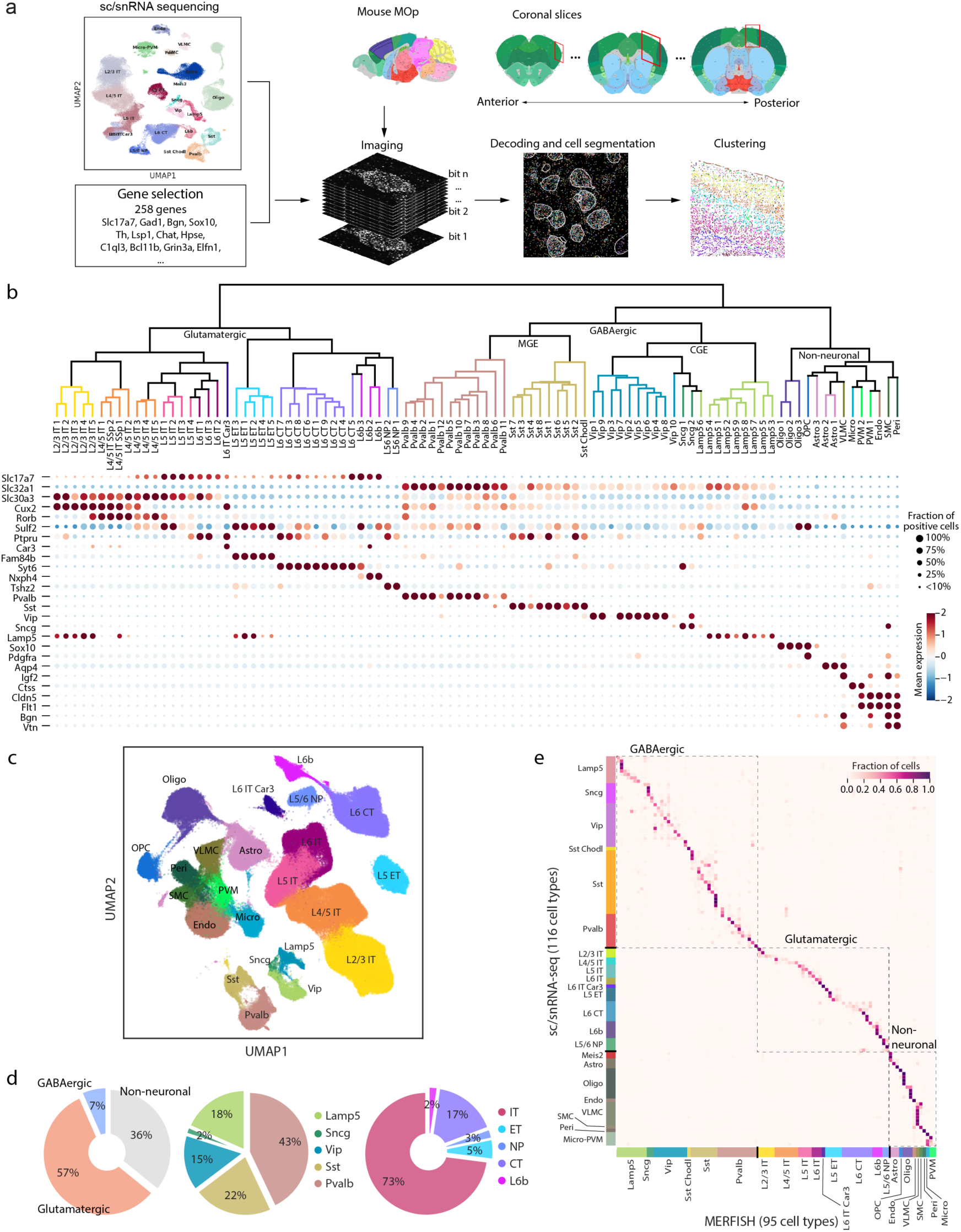
Cell type profiling and mapping of the mouse MOp by MERFISH. **a**, Schematics of MERFISH workflow for MOp cell type profiling and mapping. Based on both prior knowledge of cell type markers and sc/snRNA-seq data, we selected the MERFISH gene panel and designed the MERFISH probe set that targets these genes. Coronal slices of 10 µm-thick that contained the MOp region were cut at 100 µm intervals using the Allen CCF v3 (http://atlas.brain-map.org/) as a reference and 258 genes were imaged in each slice. The images were decoded to identify RNA species and total polyadenylated mRNA and DAPI co-stains were used for cell boundary segmentation. The decoded RNA molecules were assigned into individual cells and the resulting single-cell gene expression profiles were used for clustering analysis. **b**, (Top) Dendrogram showing the hierarchical relationship among the 39 glutamatergic, 42 GABAergic, and 14 non-neuronal clusters identified by MERFISH, constructed based on the z-scored mean cluster expressions and colored by the subclass that each cluster belongs to. (Bottom) Expression of markers genes for each subclass. **c**, UMAP of cells measured by MERFISH colored based on cell subclasses. **d**, Fraction of cells in each of the major cell classes (glutamatergic, GABAergic, and non-neuronal) (left), each of GABAergic subclasses (middle), and glutamatergic subclasses (right). **e**, Correspondence between the clusters determined by MERFISH and the consensus clusters determined by seven sc/snRNA-seq datasets. A neural-net classifier was trained on the z-scored expression profiles in the MERFISH dataset, and used to predict a MERFISH cluster label for each cell in the snRNA-seq 10x v3 B dataset generated in a companion study^38^. Cells were grouped based on their sc/snRNA-seq cluster identity, and the fraction of cells from a given sc/snRNA-seq cluster that were predicted to have each MERFISH cluster label were plotted.

We then performed MERFISH measurements on a series of coronal slices of the adult mouse brain (Bregma +2.4 to -0.6) at 100 µm intervals along the anterior-posterior axis and imaged the MOp region selected according to the Allen Mouse Common Coordinate Framework version 3 (CCF v3)^41^ (Figure 1a). Individual RNA molecules were clearly detected and identified in the tissue slices (Extended Data Figure 1a). The decoded RNA spots were assigned into individual cells segmented using 4’,6-diamidino-2-phenylindole (DAPI) and total mRNA staining (Extended Data Figure 1b). The mean copy number per cell for individual genes obtained from MERFISH were highly reproducible between biological replicates (Extended Data Figure 1c) and exhibited high correlation with the gene expression level measured by bulk RNA sequencing (Extended Data Figure 1d).

In total, we imaged and segmented ∼300,000 individual cells in the MOp in 64 coronal slices from two adult mice. Based on the single-cell expression profiles, we identified transcriptionally distinct cell populations using an unsupervised, community-detection-based clustering algorithm^42-45^ (Figure 1a). The clustering identified 39 excitatory neuronal populations, 42 inhibitory neuronal populations, and 14 non-neuronal cell populations in the MOp, as well as four distinct cell clusters exclusively outside of the MOp (in striatum or lateral ventricle) which were not included in subsequent analyses. The MERFISH-derived MOp cell taxonomy showed a hierarchical structure of multiple levels (Figure 1b). The first level of separation occurred between glutamatergic, GABAergic, and non-neuronal cell classes. The GABAergic cell class consists of two groups corresponding to their developmental origins: the CGE- and MGE-derived cells. Based on the expression of marker genes, we defined five subclasses of GABAergic cells: Pvalb, Sst, Vip, Sncg and Lamp5 (Figure 1b, bottom). The glutamatergic neuronal clusters could be grouped into the following subclasses with distinct projection properties (identified based on known marker genes^23^): layer 5 extratelencephalic projecting neurons (L5 ET, also known as pyramidal tract neurons, marked by *Fam84b*), layer 5/6 near-projecting neurons (L5/6 NP, marked by *Tshz2*), layer 6 CT neurons (L6 CT, marked by *Syt6*), layer 6b neurons (L6b, marked by *Nxph4*), as well as IT neurons (marked by *Slc30a3*), which were further divided into several subclasses based on different cortical layer assignment (L2/3 IT, L4/5 IT, L5 IT, L6 IT, plus a distinct L6 IT Car3 type). We also identified major non-neuronal cell types in MOp (Figure 1b), including astrocytes (astro), endothelial cells (endo), microglia (micro), oligodendrocyte precursor cells (OPCs), mature oligodendrocytes (oligo), perivascular macrophages (PVM), pericytes (peri), smooth muscle cells (SMC) and vascular leptomeningeal cells (VLMC). The cell subclasses determined by MERFISH (Figure 1c) showed excellent correspondence to those determined using sc/snRNA-seq in the companion paper^38^ (Extended Data Figure 2a).

Because MERFISH allows in situ cell type identification in intact tissue slices without dissociation-induced cell loss, this allowed us to determine quantitatively the composition of cells in the MOp and its vicinity. We found that the MOp was made of 57% glutamatergic, 7% GABAergic, and 36% non-neuronal cells (Figure 1d). The GABAergic cells were comprised of 43% Pvalb cells, 22% Sst cells, 18% Lamp5 cells, 15% Vip cells and 2% Sncg cells, and the glutamatergic cells were comprised of 73% IT cells, 17% CT cells, 5% ET cells, 3% NP cells and 2% L6b cells.

MERFISH data further divided the 23 subclasses of cells into 95 clusters, for which we used nomenclature style of adding a numerical index following the subclass name (e.g. Pvalb 1, L5 IT 3) (Figure 1b). Our MERFISH images also covered part of the secondary motor region (MOs) and primary somatosensory region (SSp), and in most cases, we did not separately label cells in the MOs or SSp unless the cluster was located primarily in these regions. The clusters identified by MERFISH show good correspondence to the clusters identified by sc/snRNA-seq measurements (Figure 1e) and by an integrated analysis of single-cell transcriptomic and epigenomic data in the companion paper^38^ (Extended Data Figure 2b). MERFISH analysis also revealed clusters not identified by the sc/snRNA-seq data, mostly in the form of refined cluster splitting, especially in the glutamatergic L2/3 IT and L4/5 IT subclasses. Likewise, some clusters identified by the sc/snRNA-seq data were not distinguished by MERFISH as separate clusters, mostly in the GABAergic cells. In addition, the non-neuronal cells were mostly only split at the subclass level in the MERFISH data because we only included 1 or 2 canonical markers for each major non-neuronal subclass in the MERFISH gene panel.

### Spatial organization of transcriptomically defined cell populations in MOp

Single-cell gene expression profiling and cell type identification in intact tissues by MERFISH allowed us to map the spatial organization of the 95 transcriptomically distinct cell populations in the MOp (Figure 2a). The layered organization of the glutamatergic subclasses, especially the IT subclasses, provided a laminar appearance for the overall cellular organization of the MOp (Figure 2a and Figure 2b, left). Unlike the IT cells, which spanned across nearly all cortical layers, the ET, NP, CT and L6b cells populated only deeper layers (Figure 2b left and Figure 2c). At the finer level, individual glutamatergic clusters adopted spatially distinct, partially overlapping distributions along the cortical depth direction or medial-lateral direction, and many glutamatergic clusters assumed narrow distributions along cortical depth direction that sub-divided cortical layers, often without discrete layer boundaries, as will be described in more details in a later section.

**Figure 2.**
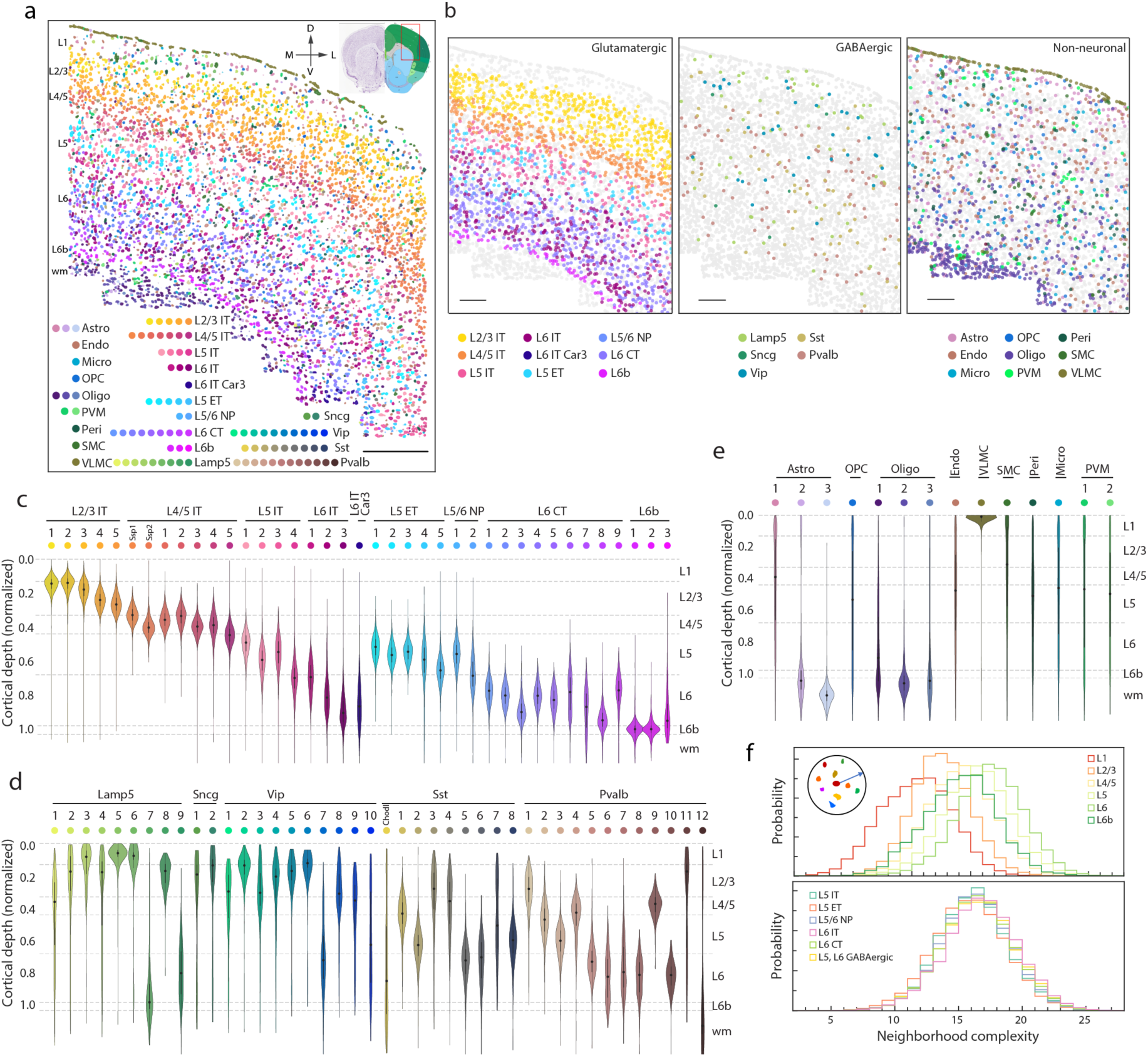
Spatial organization of cell classes, subclasses and clusters in the MOp. **a**, Spatial distribution of the 95 cell clusters determined by MERFISH shown in one of the coronal slices (Bregma ∼+1.1). Cells are shown as segmented boundaries and colored by their cluster identities. The slice is orientated as shown by the arrows. wm: white matter; M, medial; L, lateral; D, dorsal; V, ventral. Scale bar: 400 µm. **b**, Spatial distribution of the cell subclasses in the glutamatergic cell class (left), GABAergic cell class (middle), and non-neuronal cell class (right) in the same slices as shown in (a). Cells are shown as circles, with indicated cells colored by subclasses and other cells shown in grey. Scale bars are 200 µm. **c**, The distribution of the glutamatergic neuronal clusters along the cortical depth axis shown in violin plots. The cortical depth of a cell is defined as the shortest distance of its soma centroid to the cortical surface measured in each slice, and was normalized by the cortical thickness with 0 representing the cortical surface and 1 representing the median depth of the L6b cells in each slice. The dashed grey lines mark the approximate layer boundaries. **d**, Normalized cortical depth distributions of the GABAergic neuronal clusters as in (c). **e**, Normalized cortical depth distribution of the non-neuronal clusters as in (c). **f**, Spatial complexity of the cell neighborhoods in the MOp. The neighborhood complexity of a cell is defined as the number of different cell clusters present within a 100 µm-radius neighborhood surrounding the given cell. Top: Probability distributions of the neighborhood complexity of any given cell of each cortical layer. Bottom: Probability distributions of the neighborhood complexity of any given cell of in different cell subclasses in the deep layers.

The GABAergic inhibitory neurons also showed a high level of molecular diversity. At the subclass level, the CGE-derived GABAergic neurons – the Lamp5, Sncg and Vip subclasses – were more populated in the upper layers, whereas the MGE-derived GABAergic neurons – the Sst and Pvalb subclasses – were more abundant in deep layers (Figure 2b middle panel and Figure 2d), which is consistent with previous findings in the mouse cortex^46,47^. Surprisingly, at the cluster level, many of the GABAergic cell clusters also showed layered distributions and preferentially reside within one or two cortical layers (Figure 2d). The Lamp5 clusters were highly enriched in L1 and L2/3, except clusters 7 and 9, which distributed broadly in the deep layers (Figure 2d). Lamp 5 and 6 were found almost exclusively in L1 (Figure 2d). Sncg cells, a rare population in the MOp, were enriched in L1 and L2/3 (Figure 2d). Most Vip clusters were present mainly in upper layers L1 and L2/3, whereas Vip clusters 7 and 10 were more widely distributed across layers. The Sst and Pvalb neurons overall show broad distribution in the deep layers, but at the cluster level, almost all Sst and Pvalb clusters displayed restricted laminar distribution with preferential distribution in one layer (Figure 2d).

Imaging the 30 coronal slices every 100 µm along the anterior-posterior axis for each animal also allowed us to determine the distributions of neuronal clusters along this direction. We found that except for several glutamatergic clusters (e.g. L2/3 IT 1 and 2, L4/5 IT 2-4, L5 IT 3, L6 IT Car3, L5 ET 4, L6 CT 1 and 7), most neuronal clusters adopted largely uniform distributions along the anterior-posterior direction (Extended Data Figure 3).

We also mapped the spatial organizations of the non-neuronal cells (Figure 2c right panel and Figure 2e). Among the three astrocyte clusters, the most abundant cluster, Astro 1, showed no preference in laminar residency, a less abundant cluster, Astro 2, showed enrichment in L1 and the white matter, and the rarest type, Astro 3, was found almost exclusively in the white matter. The oligodendrocyte lineage was divided into OPCs and three mature oligodendrocyte clusters. The mature oligodendrocytes were enriched in the white matter, accounting for ∼90% of the cells in the corpus callosum, whereas the OPCs distributed evenly across all layers. The VLMCs formed the outmost layer of cells of the cortex. The other non-neuronal cell types, including microglia, PVMs, SMCs, pericytes and endothelial cells, exhibited more disperse distributions across the cortical layers.

We noticed substantial spatial intermixing of different cell populations across the MOp, with individual neighborhoods adopting a complex cell composition. To quantify the complexity of the cell composition in the neighborhood of each cell, we determined the number of distinct cell clusters that were present in the 100 µm vicinity of each cell and determined the distributions of this complexity metric for cells in each cortical layer and in each subclass (Figure 2f and Extended Data Figure 4). For each cell neighborhood, we observed a large number of cell clusters, indicating a high level of local cellular heterogeneity, and the composition complexity of cell neighborhood increased as the cell soma moved towards deeper layers (Figure 2f top). In the deep layers L5 and L6, where the cell neighborhood was the most complex, different cell types exhibited comparable degree of neighborhood complexity (Figure 2f bottom).

### Diversity of L5 ET, L5/6 NP, L6 CT and L6b neurons

Transcriptomically, the L5 ET, L5/6 NP, L6 CT and L6b subclasses of neurons appeared as discrete cell populations (Figure 3a). Each subclass was subdivided into finer clusters (Figure 3a, Extended Data Figure 5). Spatially, the five L5 ET clusters were segregated into 2 sublayers with the L5 ET 5 cluster distributed in the lower part of layer 5 and the L5 ET 1-3 clusters intermixed in upper layer 5 (Figure 3b). L5 ET 4 located at the lateral side of MOp (Figure 3b) and was found to be more abundant in the anterior part of MOp whereas in the posterior part, L5 ET 4 mostly resided outside of MOp (Extended Data Figure 3). It has been previously reported that two distinct L5 ET populations in upper and lower layer 5 of the anterior lateral motor cortex (ALM) project to thalamus and medulla, respectively, and played specialized roles in motor control in the ALM^48^. In a companion BICCN paper, Zhang et al also identified a distinct medulla-projecting ET sub-type and several other ET sub-types with less projection specificity using the epi-retro-seq method^49^. The lower layer 5 cluster L5 ET 5 identified here by MERFISH corresponded to the L5 ET (1) cluster identified by integrated analysis of single-cell transcriptomic and epigenomic data in the companion paper^38^ (Extended Data Figure 5), which in turn corresponds to the medulla-projecting ET cluster identified by epi-retro-seq^49^. Our MERFISH data further showed that this unique medulla-projecting L5 ET 5 cell type was mainly present in MOp but rarely in the adjacent SSp region (Extended Data Figure 6). The L5/6 NP neurons were divided into two clusters (Figure 3a), with L5/6 NP 1 found mainly in layer 5 and L5/6 NP 2 found in deeper regions extending into layer 6 (Figure 3c).

**Figure 3.**
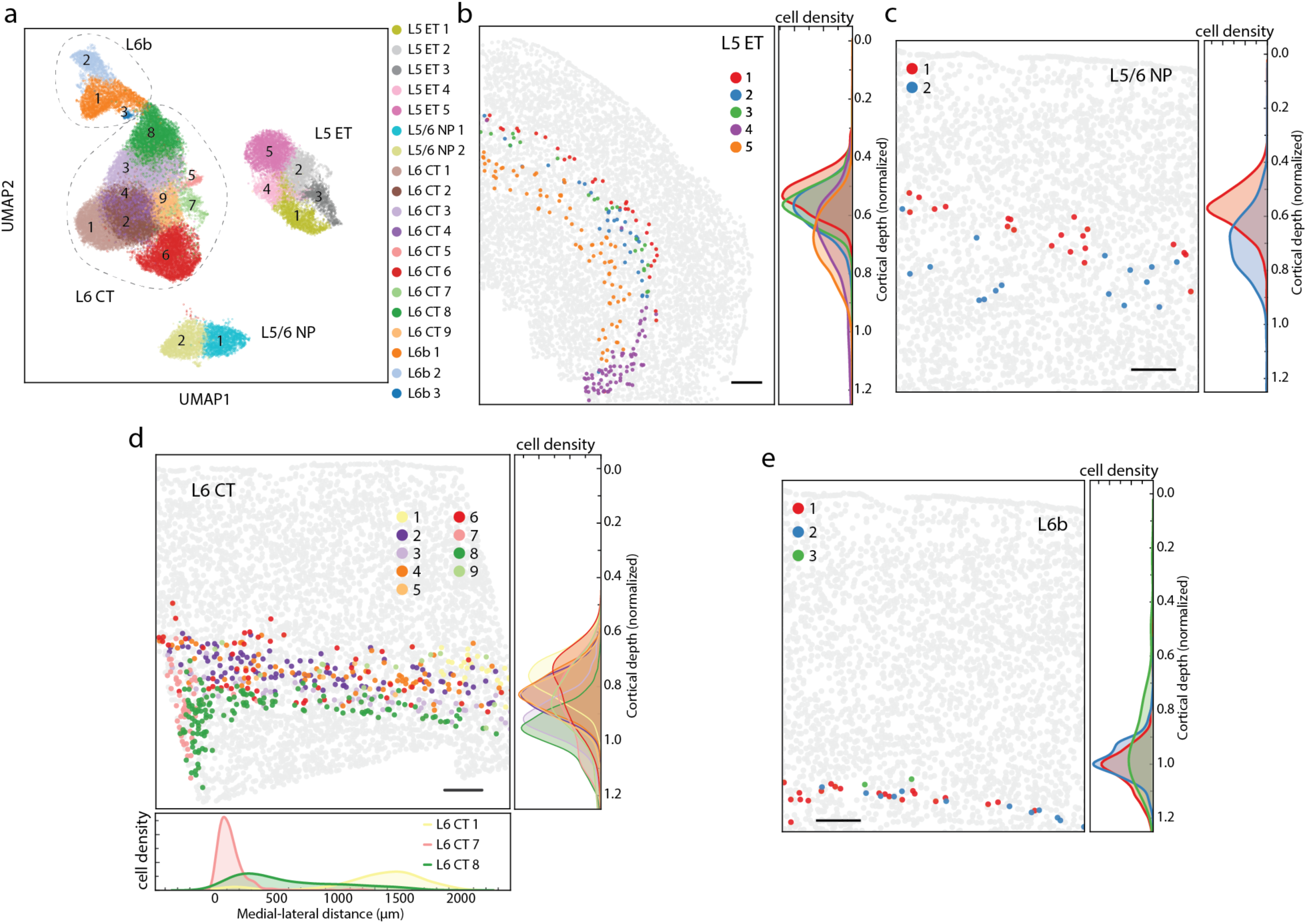
Diversity and spatial organization of the ET, CT, NP and L6b neurons. **a**, UMAP of the ET, CT, NP and L6b neurons colored by the MERFISH cluster identity. **b**, Spatial distribution of five L5 ET cell clusters in one of the coronal slices (∼Bregma +1.4). L5 ET cells are highlighted in colors and other cells are shown in grey. Normalized cortical depth distributions of the cells of each L5 ET cluster is shown in the right panel. **c**, Spatial distribution of two L5/6 NP cell clusters in one of the coronal slices (∼Bregma +0.4), as in (b). **d**, Spatial distribution of the nine L6 CT cell clusters in one of the coronal slices (∼Bregma +0.3), as in (b). The medial-lateral (ML) distribution for clusters 1, 7 and 8 is additionally shown at the bottom. The ML distribution is calculated in each coronal slice in which cluster 7 is present, where the ML distance of zero is set to be the location of the cluster 7 cell that is the closest to the midline. **e**, Spatial distribution of the three L6b clusters in one of the coronal slices (∼Bregma +0.4), as in (b). Scale bars in (b-e) are 200 µm.

The L6 CT neurons, the most abundant subclass in L6, were divided into nine clusters (Figure 3a). These clusters exhibited a complex spatial pattern, with distinctions both in the cortical depth direction and in the medial-lateral direction (Figure 3d). The L6 CT 1 cluster located preferentially at the lateral side of MOp, which likely represents an extension of a CT type from the SSp region. The L6 CT 7 cluster, on the other hand, was found exclusively at the medial side of MOp in the posterior part, which is close to the MOs region and the anterior cingulate area. Clusters L6 CT 6, 4, 3 and 8 spanned the entire medial-lateral range of the MOp and exhibited a layered organization from top to bottom of layer 6. L6b cells, which formed the innermost layer of the neocortex, were subdivided into three clusters (Figure 3a), but these three clusters were completely intermixed in space, showing no difference in spatial distribution along the anterior-posterior axis, medial-lateral axis, or cortical depth axis (Figure 3e).

Taking advantage of the correspondence between MERFISH clusters and the clusters identified by integrated analysis in the companion paper^38^ (Extended Data Figure 5), we can further infer addition information, such as the epigenomic signatures, about the spatially-resolved MERFISH clusters. For example, *Foxp2*, a marker gene identified for L6 CT 1 and 2 MERFISH clusters, showed enrichment of open chromatin reads and reduced DNA methylation in the corresponding integrated cluster L6 CT Cpa6 (1), as compared to the L6 CT Cpa6 (4) and Nxph2 Kit clusters, which correspond MERFISH clusters L6 CT 8 and 7, respectively (Extended Data Figure 7). Based on the correspondence between MERFISH and sc/snRNA-seq or integrated clusters, or more sophisticated integration analysis, it is also possible to impute marker genes for MERFISH clusters that were not included in the MERFISH gene panels, which would not only provide a more complete marker gene sets for the MERFISH clusters, but also allow the prediction of spatial expression patterns for these gene.

### A continuous spectrum of the IT neurons

The IT neurons constitute the largest branch of neurons in the MOp, which span nearly the entire MOp region from L2/3 to L6. Our MERFISH data divided the IT cell into 20 clusters, 19 of which belonged to 4 subclasses classified by the cortical layers (L2/3 IT, L4/5 IT, L5 IT and L6 IT) (Figure 4a, b). The remaining one formed a distinct cell type (Figure 4a), which corresponded to the L6 IT Car3 type identified by sc/snRNA-seq and integrated analysis (Extended Data Figure 8a and b). These L6 IT Car3 cells did not form a laminar distribution, but were located at the lateral edge of MOp (Extended Data Figure 9a).

**Figure 4.**
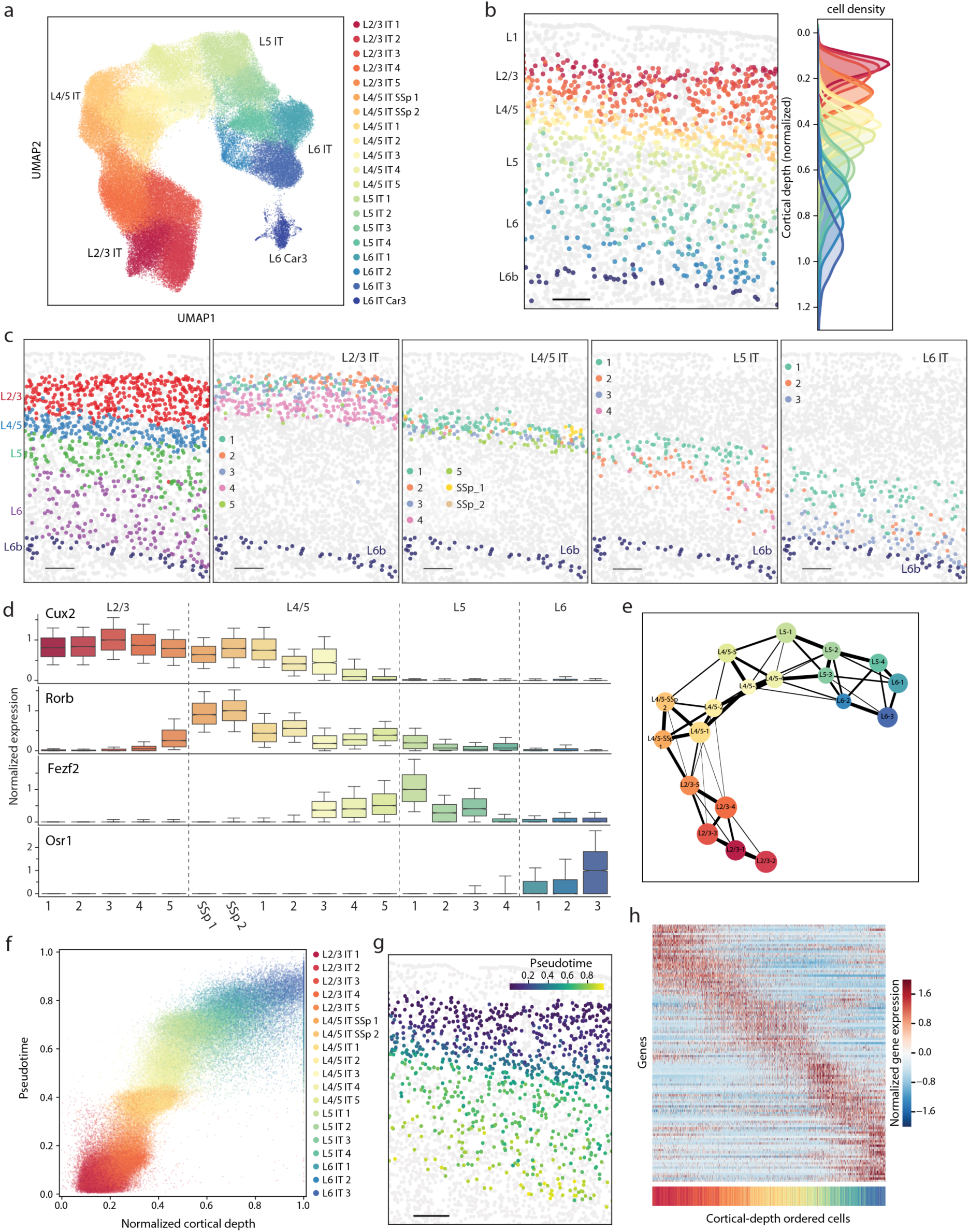
A continuous spectrum of IT cell with gene expression profile correlated with cortical depth position. **a**, UMAP of the 20 IT neuronal clusters. **b**, The IT neurons in one of the coronal slices (∼Bregma +0.4). The IT neurons are colored by their cluster identity as in (a) shown together with L6b cells in dark blue to mark the bottom border of the cortex, and all other cells are shown in grey. Scale bar: 200 µm. **c**, Spatial distribution of the IT subclasses (L2/3, L4/5, L5, and L6, left panel) and individual clusters in each subclass (right panels) shown in one of the coronal slices (∼Bregma +1.0). The L6b cells are shown in dark blue to mark the bottom border of the cortex. Scale bars: 200 µm. **d**, Box plot of the expression of layer specific marker genes (*Cux2, Rorb, Fezf2, Osr1*) in each IT neuronal cluster normalized to the maximum median expression across all clusters. The box represents the interquartile range, the median is indicated with a line, notch represents ± 95% confidence interval, whiskers are 10th and 90th percentile, outliers are not shown. **e**, Force-directed graph representing the degree of connectivity between clusters in a k-nearest neighbor (kNN) graph for the IT neuronal clusters. The expression profile of the IT cells was z-score normalized; principle-component analysis was used to reduce dimensionality to the first 19 principle components then input to construct a kNN graph. Partition-based graph abstraction (PAGA) was used to compute the connectivity between clusters in the kNN graph^53^, which is plotted with each cluster represented as a node, and the weighted edges between nodes representing their connectivity. Edges with weights below 0.1 were discarded. **f**, Scatter plot of the pseudotime vs. normalized cortical depth for individual IT neurons colored by the cell clusters. Pseudotime was calculated using an L2/3 IT 1 cell as the root, using the same kNN graph as described in (e). **g**, The same tissue slice as shown in (b) with the IT neurons colored by their pseudotime. **h**, Normalized expression of 118 differentially expressed genes of all the IT neurons across cortical depth. All IT cells were sorted in the order of ascending cortical depth and the genes are sorted by the cortical depth at which they exhibit maximal expression. The colored bar beneath the heatmap indicates the cluster identity of the cell in that column of the heatmap.

In contrast, the 19 clusters in the L2/3, L4/5, L5 and L6 IT subclasses showed fine laminar organizations (Figure 4b). These four subclasses were each subdivided into several clusters, further parcellating each layer into finer sub-layers, but without discrete boundaries (Figure 4a-c). Notably, MERFISH identified seven IT clusters residing between L2/3 and L5, which we named L4/5 IT (Figure 4c). Among these, L4/5 IT SSp 1 and 2 were located in the neighboring SSp region and partially extended into the MOp (Extended data Figure 9b), whereas the other five clusters (L4/5 IT 1-5) were located within the MOp. The MOp has been traditionally considered lacking a distinct layer 4 due to the absence of clear cytoarchitecture features^50^. Our results suggest the presence of layer 4 neurons in the MOp, corroborating results from joint analyses of multiple different experimental modalities in the BICCN consortium (see the BICCN flagship paper) and a recent report of L4-like neurons based on their anatomic and connectivity properties^51^.

Along the cortical depth direction, individual IT clusters partially overlapped in space with adjacent clusters and there was no clear difference in the extent of spatial overlap when the adjacent clusters were from the same or a different subclass (Figure 4b). Moreover, the differentially expressed genes often marked multiple cell clusters across different subclasses and exhibited gradual changes along the cortical depth direction, as represented by the cortical layer markers *Cux2, Rorb, Fezf2* and *Osr1* (Figure 4d). The two-dimension visualization by Uniform Manifold Approximation and Projection (UMAP)^52^ also revealed a gradual transition of the gene expression profiles among the IT cell clusters, except for L6 IT Car3 (Figure 4a). The apparent lack of discreteness among the IT cell clusters in both their gene expression profiles and spatial profiles led us to evaluate whether the IT cell clusters reflect cross-sections of a continuous spatial and molecular landscape.

To this end, we first quantified the similarity in gene expression between pairs of IT clusters by computing the degree of inter-cluster connectivity in a k-nearest neighbor (kNN) graph constructed based on the gene expression profiles of individual IT cells^53^. This analysis showed that the IT cell clusters formed a single, interconnected network with clusters exhibiting the highest connectivity (i.e. similarity in gene expression) to those that were spatially adjacent (Figure 4e). Next, we applied pseudotime analysis to order the IT cells based on their expression profiles. Pseudotime analysis has been most commonly used to understand dynamic or developmental states of cells^54^, but here we use this analysis to evaluate how an expression-defined trajectory of IT cells relate to their spatially-defined trajectory. Notably, ordering of cells in pseudotime was highly correlated with their cortical depth positions (Figure 4f, g and Extended data Figure 10a), and individual cells formed a continuous cloud, instead of discrete clusters, along the pseudotime and cortical depth axes (Figure 4f). We further identified genes of which the expression changed substantially with cortical depth, yielding a list of 118 genes that included the well-known layer markers as shown in Figure 4d. Sorting these genes based on the cortical depth at which they exhibited maximum expression revealed a largely gradual change of gene expression profiles of cells along the cortical depth axis, with somewhat sharper changes at the cortical depths that approximately separate subclasses (Figure 4h). Similar patterns were also observed when cells and genes were sorted based on pseudotime instead of cortical depth (Extended Data Figure 10b). At the individual gene level, the expression profile of the variable genes changed gradually along the cortical depth direction (Extended Data Figure 10c)

Taken together, these results show that the IT cells do not follow the rule of cortical layer delineation but instead adopt a largely continuous gradient distribution across the cortical depth, with the gene expression profiles of individual cells being highly correlated with their cortical depth positions.

### Complex projection patterns of the IT neurons

The axonal projection properties of the neurons are key to connect their molecular profiles with function. We thus sought to integrate MERFISH with retrograde tracing to simultaneously determine the expression profiles and spatial organization of cell types in the MOp and their projection targets at the single cell level.

To this end, we injected a retrograde tracer, cholera toxin subunit b (CTb), into three cortical regions, MOs, SSp, and the temporal association area (TEa), all of which have been reported to receive inputs from the MOp^8,9^. We injected three distinct dye-conjugated CTbs (CTb-AlexaFluor 647, 555, and 488) into MOs, SSp and TEa, respectively, and identified the cells in the MOp that projected to these injection sites by imaging the signals of the CTb conjugates. We then imaged the 258-gene MERFISH panel, as described above, in the same tissue slices for cell type identification (Figure 5a, b). This allowed individual cells to be assigned with both a cell type identity and a projecting target (Figure 5c). We performed this analysis for the MOp upper limb region (Bregma +0.7 to +0.1), and observed that ∼90% of MOs-, SSp- and TEa-projecting neurons were IT neurons and L6b neurons. Spatially, the MOs-projecting and SSp-projecting neurons were enriched in upper layers L2/3 and L4/5; the TEa-projecting neurons showed enrichment in upper L2/3, deep L4/5 and L6 (Figure 5d), consistent with previous observations^9^.

**Figure 5.**
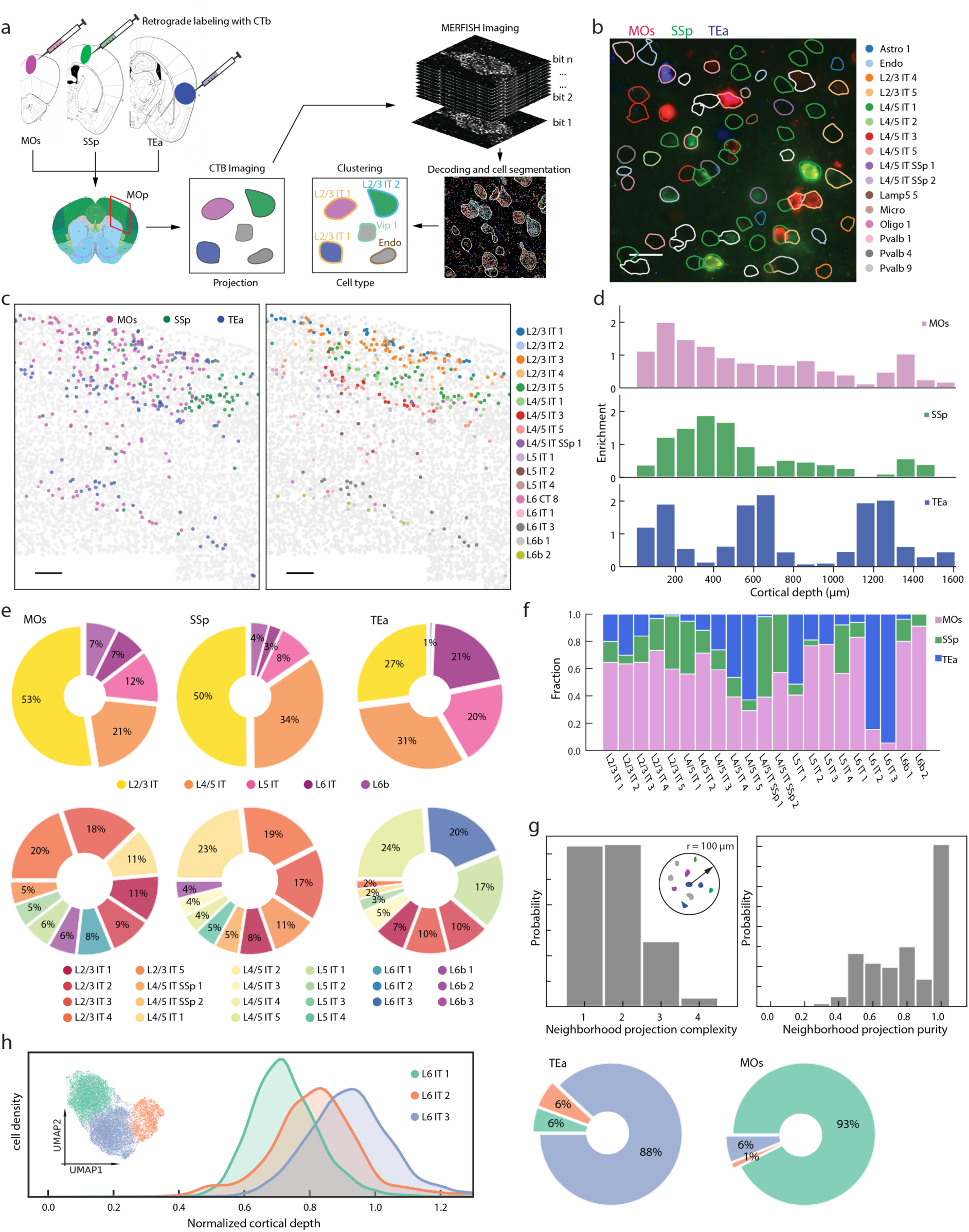
Projection patterns of the IT neurons determined by the integration of MERFISH with retrograde tracing. **a**, Schematics of workflow integrating retrograde tracing and MERFISH. Three cholera toxin subunit b (CTb) conjugates, CTb-AlexaFluor647, CTb-AlexaFluor555, and CTb-AlexaFluor488, were injected into three different cortical regions, MOs, SSp, and TEa, respectively. The coronal slices containing the MOp region were then collected and the retrograde CTb labels and MERFISH gene panel were imaged. **b**, CTb fluorescence image of a single field-of-view in the MOp with CTb-AlexaFluor647, CTb-AlexaFluor555, and CTb-AlexaFluor488 signals shown in red, green and blue, respectively. The cells boundaries were colored by their cell cluster identities determined by MERFISH. **c**, A coronal slice (∼Bregma +0.1) highlighting the CTb-positve cells colored by the injection sites (Red: MOs, Green: SSp, Blue: TEa; left) and by their cell cluster identities (right). Only the cell clusters with 3 or more cells labeled by CTb are highlighted. Scale bars: 200 µm. **d**, Histogram showing enrichment of MOs-projecting, SSP-projecting and TEa-projecting cells at different cortical depths. Enrichment was determined by comparing the fraction of relevant CTb-positive cells in each cortical-depth bin with the fraction of all IT and L6b cells in the same bin. **e**, Left: Pie charts showing the proportions of MOs-projecting cells belonging to each cell subclasses (top row) and cell clusters (bottom row). Only top 10 cell clusters in each case are shown in the bottom row. Middle: same as left but for SSP-projecting cells. Right: same as left but TEa-projecting cells. **f**, Fractions of MOs-projecting, SSP-projecting and TEa-projecting cells in each cell cluster. **g**, Neighborhood projection complexity (left) and purity (right) of the cells in the MOp. The neighborhood projection complexity of a CTb-positive cell is defined as the number of different projection targets (MOs, SSp, TEa, or CTb-negative) of the cells belong to the same cluster present within a 100 µm-radius neighborhood surrounding the cell. Neighborhood projection purity of a CTb-positive cell is defined as the fraction of all cells of the same cluster within the 100 µm-radius neighborhood that belong to the most abundant projection target. The normalized histograms of the neighborhood projection complexity and purity for all CTb-positive cells are shown. **(h)** The projection specificity of the molecularly and spatially similar L6 IT clusters. The cortical depth distributions (left) and UMAP (left inset) of L6 IT 1, 2 and 3 indicate the similarity in gene expression profiles and spatial locations between the three clusters. Pie charts (right) show the relative proportions of the three clusters that projected to MOs and TEa, respectively.

At the single cell and cell cluster level, we observed a complex projection pattern that connected MOp IT neurons and the three target regions. Notably, even neighboring IT neurons in the same cell clusters could send output to different target regions, and likewise, the same target region could also receive inputs from different cell clusters (Figure 5c, e). At the subclass level, MOs and SSp received inputs from all five subclasses of IT neurons and L6b neurons but both were dominated by L2/3 and L4/5 IT neurons, and TEa received input from the four IT subclasses (L2/3, L4/5, L5 and L6) with nearly equal contributions but had almost no input from L6b neurons (Figure 5e, upper panel). At the cluster level, all three regions received inputs from a large number of individual clusters, each region from a quantitatively different composition of clusters (Figure 5e, lower panel).

Out of the 22 IT and L6b cell clusters, 21 clusters showed projection to the MOs, SSp and TEa regions. All of these clusters targeted more than one region (Figure 5f). Even spatially adjacent cells belonging to the same clusters could project to different regions (Figure 5g). These results suggest that the projection target of a neuron is not simply dictated by its gene expression profile and spatial location.

Interestingly, although IT neurons exhibited a gradual change in their gene expression profiles and spatial distributions, some of them showed highly specific projection patterns. For example, almost all CTb-positive L6 IT 3 neurons targeted the TEa but not MOs and SSp, whereas the vast majority of the CTb-labeled L6 IT 1 neurons targeted MOs but not SSp and TEa (Figure 5f). Conversely, among the L6 IT neurons, MOs almost exclusively received input from the L6 IT 1 cluster and TEa almost exclusively received input from the L6 IT 3 cluster, despite the similar expression files and substantially overlapping spatial distributions of these cells (Figure 5h). How such discrete and specific projection property arises from these neurons with similar and gradually varying expression profiles and spatial distributions, whether this is due to a molecular signature not capture by transcriptomic profiling or has arisen from a developmental origin, remains to be determined.

## Discussion

In this study, we used MERFISH to generate a molecularly defined and spatially resolved map of cell populations for the mouse MOp. The cell census defined by MERFISH, including 95 neuronal and non-neuronal populations, showed good correspondence to that defined by the single-cell sequencing-based transcriptomics datasets reported in a companion BICCN paper^38^. The MOp cell census determined by these approaches showed both similarity and differences to the cell census of other cortical regions^23,38,55^.

The direct spatial measurements by MERFISH allowed us to map the spatial organization of the 95 molecularly defined cell clusters with high resolution. Our results showed laminar restrictions for different subclasses of neurons that are consistent with previous findings^22,23,46,47^, but also revealed a previously unknown, high-resolution spatial map for individual neuronal clusters. We observed that most transcriptomically distinct cell clusters adopted distinct spatial distributions. Notably, our MERFISH images revealed a laminar organization of not only excitatory neurons, but also inhibitory neurons, with many inhibitory neuronal clusters preferentially located in one or two cortical layers. Moreover, many excitatory neuronal clusters adopted narrow distributions along the cortical depth direction that subdivided individual cortical layers into finer laminar structures, often without discrete layer boundaries.

We noticed that although neurons tended to form discrete populations of cells with distinct expression profiles at the subclass level, the clusters within individual subclasses often exhibited more gradual changes, adding evidence to the co-existence of discrete and continuous cell heterogeneity^23,56,57^. Remarkably, our results showed that the entire cohort of IT cells (barring the small Car3 cluster), which consist of several subclasses and constitute >70% of all excitatory neurons in the MOp, formed a largely continuous spectrum of cells instead of discrete clusters. Continuous variations in gene expression was also observed among IT cells in the isocortex by an concurrent, independent scRNA-seq study^55^. Here, with spatially resolved single-cell gene-expression profiling afforded by MERFISH, we observed concurrent gradual changes among IT neurons both in gene expression space and in real space, with a strong correlation between the expression profiles of individual neurons and their cortical depth positions, revealing a continuous molecular and spatial gradient of cells spanning nearly the entire cortical depth.

Taken together, our MERFISH measurements of the ∼100 neuronal and non-neuronal cell populations revealed the correspondence between molecular diversity and spatial organization of cells in the cortex with an unprecedented resolution and granularity.

We further investigated how individual molecularly identified cell types correlate with their projection targets by integrating MERFISH with retrograde tracing. This approach allowed in situ determination of both gene expression profiles and projection targets with single cell resolution and hence enabled the determination of source neurons projecting to specific target regions with high spatial and molecular resolution. Our results showed that projections of MOp neurons to other cortical regions did not occur in a one-cell-type-to-one-target-region manner, but rather formed a complex many-to-many network: each cell cluster can project to multiple regions and each region can receive input from many clusters, with an underlying specificity. Here, our proof-of-principle measurements probed only three target regions, but more target regions could be measured to allow the construction of a more compressive projection map for the MOp cell types. Technologically, the throughput of projection targets profiled per experiment could also potentially be scaled up by using different projection-labeling approaches, such as oligonucleotide-labeling of CTb or high-diversity barcoded tracers^58^, to allow many distinct tracers to be imaged and hence many target regions to be interrogated in each MERFISH experiment. Ultimately, we envision that MERFISH may also be combined with barcoded, trans-synaptic viral tracers to allow the generation of a high-resolution cell-type-to-cell-type connectivity map.

In this BICCN consortium, a multitude of experimental modalities, including single-cell transcriptomics, single-cell epigenomics, anatomical measurements, and electrophysiology have been used to provide cell census and atlas of the MOp (see companion papers^38,49,59-61^). Some of the studies probed two or three properties of the neurons simultaneously. In addition to our study, which combined MERFISH and neural tracing to determine the gene expression profiles, spatial locations, and projection targets of the same neurons, the companion epi-retro-seq paper^49^ simultaneously probed the epigenomic properties and projection targets and the companion patch-seq study^60^ jointly probed morphological, electrophysiological and transcriptomic properties of individual neurons. In addition to the transcriptomic profiles, many other properties can also be probed by imaging-based methods, including epigenomic properties (e.g. chromatin conformation^62,63^ and accessibility^64^), anatomic properties (e.g. cell morphology and projection), as well as functional properties (e.g. by using neuronal activity markers). Imaging-based methods are often intrinsically compatible, and can be performed in a manner that preserves the sample for other analyses. We thus envision an exciting possibility of combining single-cell transcriptome imaging with all these other imaging-based approaches to simultaneously probe the molecular, spatial, anatomical and functional properties of individual cells, expediting our understanding of how different cell types are connected to form functional circuits in the brain.

## Data and code availability

All raw and processed MERFISH data can be accessed via the Brain Image Library (BIL) ftp archive: ftp://download.brainimagelibrary.org:8811/02/26/02265ddb0dae51de/. Code for MERFISH image analysis is available at https://github.com/ZhuangLab/MERlin.

## Supplementary Information

is linked to the online version of the paper.

## Acknowledgments

This work was supported in part by the National Institutes of Health (U19MH114830 and U19MH114821). X.Z. is a Howard Hughes Medical Institute Investigator.

## Author Contributions

M.Z., S.E., B.Z., Z.Y., H.Z., H.D. and X.Z designed the experiments. M.Z. and Z.Y. designed the MEFISH gene panel. M.Z. performed MERFISH experiments. B.Z. performed CTb tracer injection. M.Z. and S.E. performed data analysis. M.Z., S.E. and X.Z. wrote the paper with input from B.Z., Z.Y., H.Z, and H.D. X.Z. oversaw the project.

## Author Information

X. Z. is a co-founder of Vizgen. Correspondence and requests for materials should be addressed to X.Z.

## Extended Data Figures

**Extended Data Figure 1.**
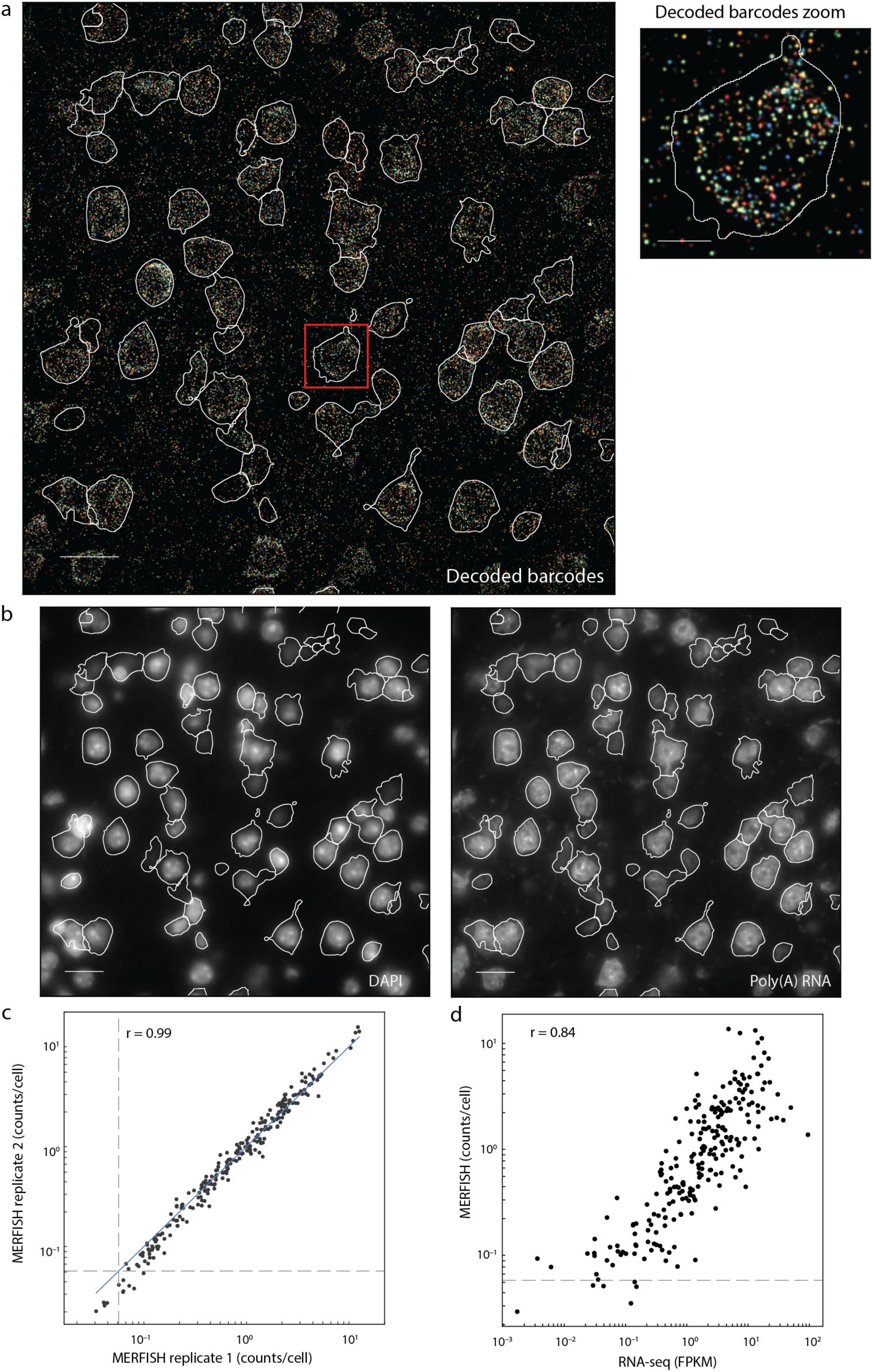
RNA identification and cell segmentation of MERFISH image, replicate reproducibility of MERFISH data, and correlation between MERFISH and bulk RNA-seq results. **a**, Decoded MERFISH image of a single field-of-view, shown as a maximum intensity projection across all seven z-planes. In these experiments, we assigned 22-bit Hamming Distance 4, Hamming Weigh 4 barcodes capable of error detection and correction to individual RNA species, physically imprinted the barcodes onto the RNAs using a high-diversity of oligonucleotide probes (encoding probes), and detected the barcodes bit-by-bit using sequential imaging. Each encoding probe comprises a target sequence that specifically binds to one of the targeted genes and two readout sequences, each readout sequence corresponds to one bit, and the presence of specific readout sequences on the entire set of encoding probes bound to an RNA determines which bits read “1” for this RNA. The 22 bits were imaged in eleven sequential rounds of hybridization with two-color imaging per round. The decoded image shows all pixels that belonged to detected correct barcodes. Each barcode was assigned a unique RGB value, and pixels were colored based on their assigned barcode. The intensity of each pixel was scaled based on the L2-norm of their fluorescence signal intensity across all bits. Right inset: The boxed region of the image shown at a greater magnification. Segmented cell boundaries as determined in (b) are shown in white. Scale bars are 20 µm and 5 µm, for the full images and the magnified region in the insets, respectively. **b**, DAPI (left) and poly(A) RNA (right) images for the same field-of-view as in (a), shown is the central z-plane (z = 4.5 µm). These images are used to define the boundaries of each cell using a seeded watershed algorithm, with DAPI defining the seed and poly(A) used to define the extent of each cell (see Online Methods). The cell boundaries are shown in white. Scale bars are 20 µm. **c**, Scatterplot of the average counts of individual genes per cell for the two biological replicates, showing all genes measured by MERFISH. The blue solid line indicate equality. The grey dashed lines indicate the average counts per cell of the blank barcodes (i.e. valid barcodes that were not assigned to any RNA), which provides an indication of the false-positive rate. The Pearson correlation coefficient is 0.99. **d**, Scatterplot of average copy number per cell of individual genes determined by MERFISH versus expression level determined by bulk RNA-seq for the MOp region for all genes measured by MERFISH. The dashed line indicates the average copy number per cell of the blank barcodes. The Pearson correlation coefficient is 0.84.

**Extended Data Figure 2.**
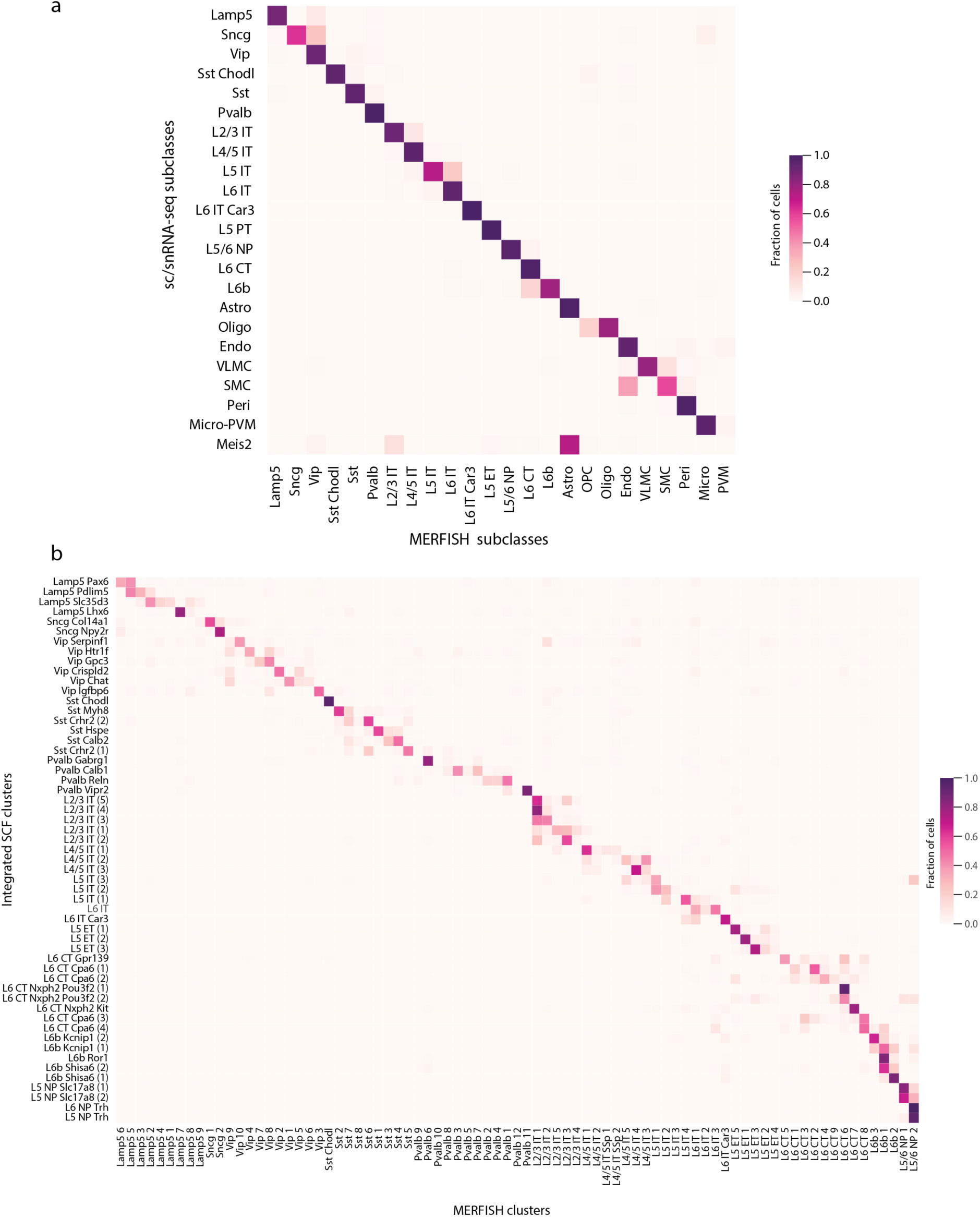
Correspondence between cell subclasses and clusters identified by MERFISH and those identified by single-cell sequencing based approaches. **a**, Correspondence between cell subclasses identified by MERFISH and by sc/snRNA-seq. A neural-net classifier was trained to predict MERFISH subclass labels using the z-scored expression profiles of individual cells in the MERFISH data. The snRNA-seq 10x v3 B dataset was z-scored, and then the subset of genes in the MERFISH gene panel were used along with the trained model to predict a MERFISH subclass label for each cell in the snRNA-seq dataset. From this, each snRNA-seq cell had both a predicted MERFISH subclass label and a subclass label determined from the consensus sc/snRNA-seq clustering^38^. Cells were grouped based on their consensus sc/snRNA-seq cluster identity, and then the fraction of cells from a given consensus sc/snRNA-seq subclass that were predicted to have each MERFISH subclass was determined and plotted. **b**, Correspondence between neuronal clusters identified by MERFISH and by integrated analysis of sc/snRNA-seq, snATAC-seq and snmC-seq. A classifier was trained as in (a), but in this case only neuronal clusters identified by the MERFISH measurements were considered, and MERFISH cluster labels were predicted for the cells in the snRNA-seq 10x v3 B dataset and compared with their cluster identities determined by the integrated clustering analysis of sc/snRNA-seq, snATAC-seq and snmC-seq measurements using SingleCellFusion (SCF)^38^ (55 out of the total 56 neuronal integrated clusters revealed by SCF is detected in the snRNA-seq 10x v3 B data used here).

**Extended Data Figure 3.**
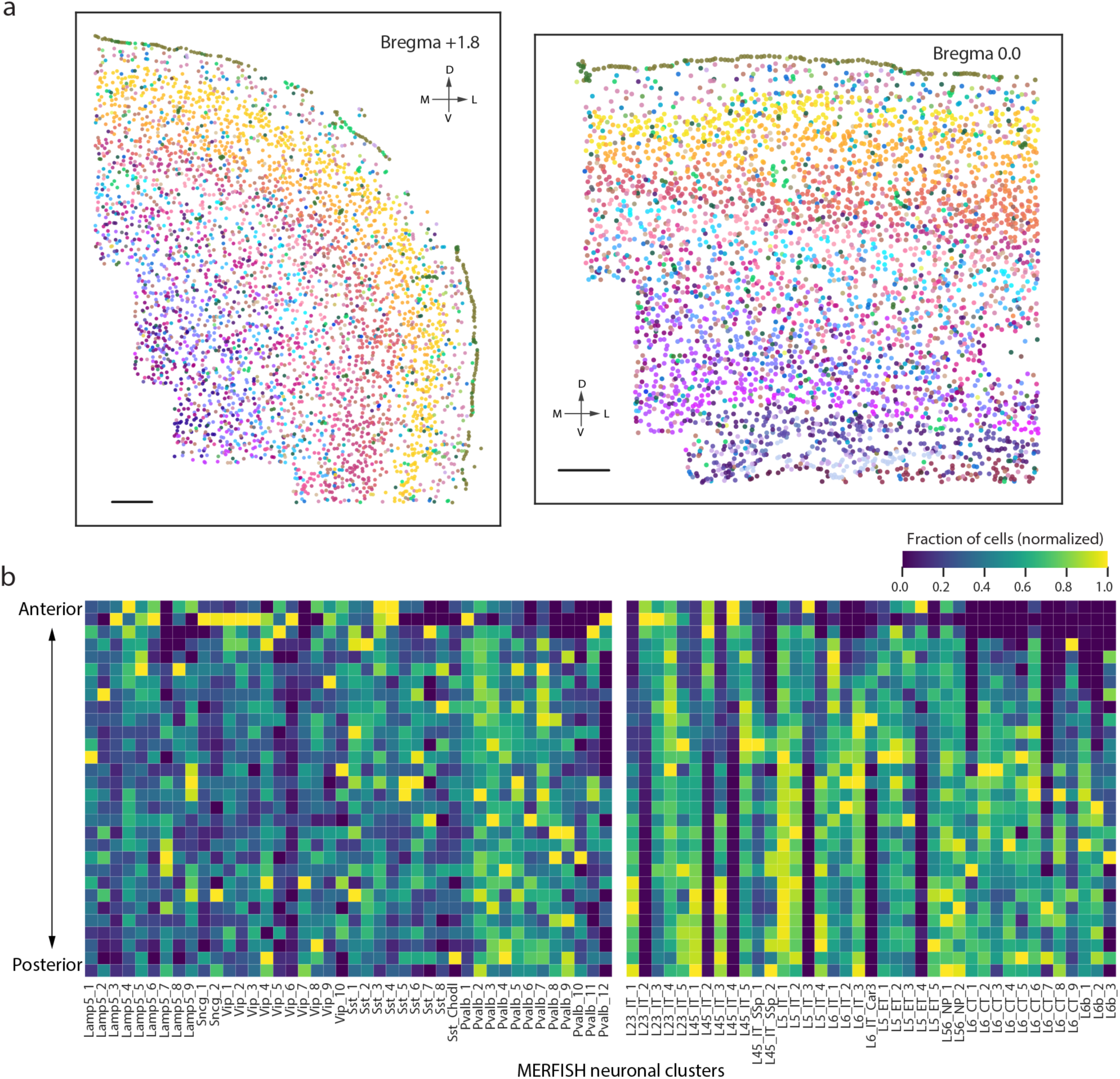
Anterior - posterior distribution of neuronal clusters in the MOp. **a**, Spatial organization of cells in two additional representative coronal sections at Brigma +1.8 and 0.0. Cells are colored by subclass, as in Figure 2a. Scale bars are 200 µm. **b**, Heatmap quantifying the anterior - posterior distribution of the neuronal clusters. Slices were arranged from anterior-most to posterior-most based on their bregma coordinates. For each cluster, the fraction of cells found in each slice was determined and normalized to the maximum across all slices.

**Extended Data Figure 4.**
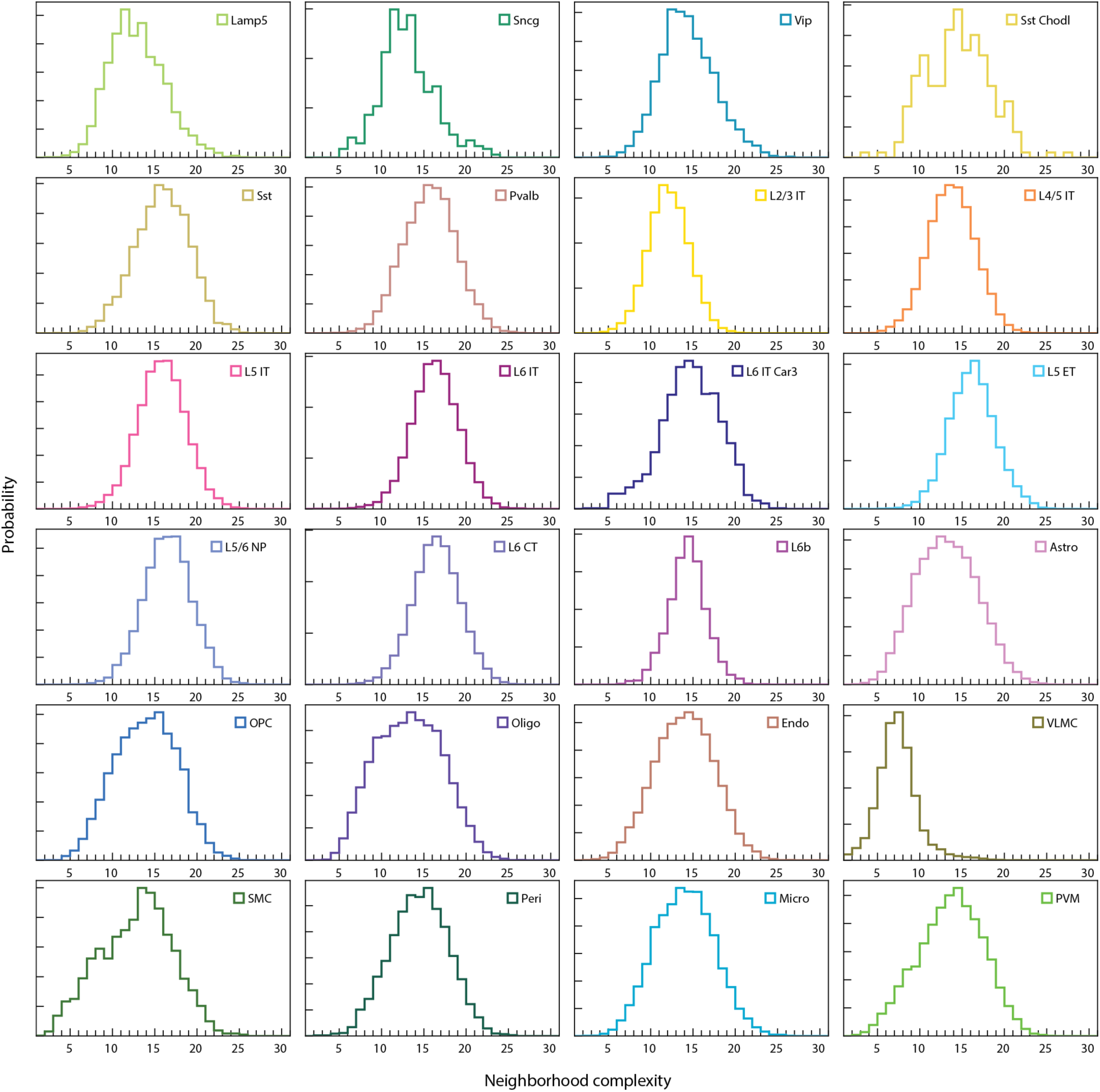
Neighborhood complexity of individual cells belonging to different subclasses. The neighborhood complexity of a cell is defined as the number of different cell clusters present within a circle of 100 µm radius around the cell of interest, as in Figure 2f. A normalized histogram of the neighborhood complexity for all cells from a given subclass is shown for each subclass identified by MERFISH.

**Extended Data Figure 5.**
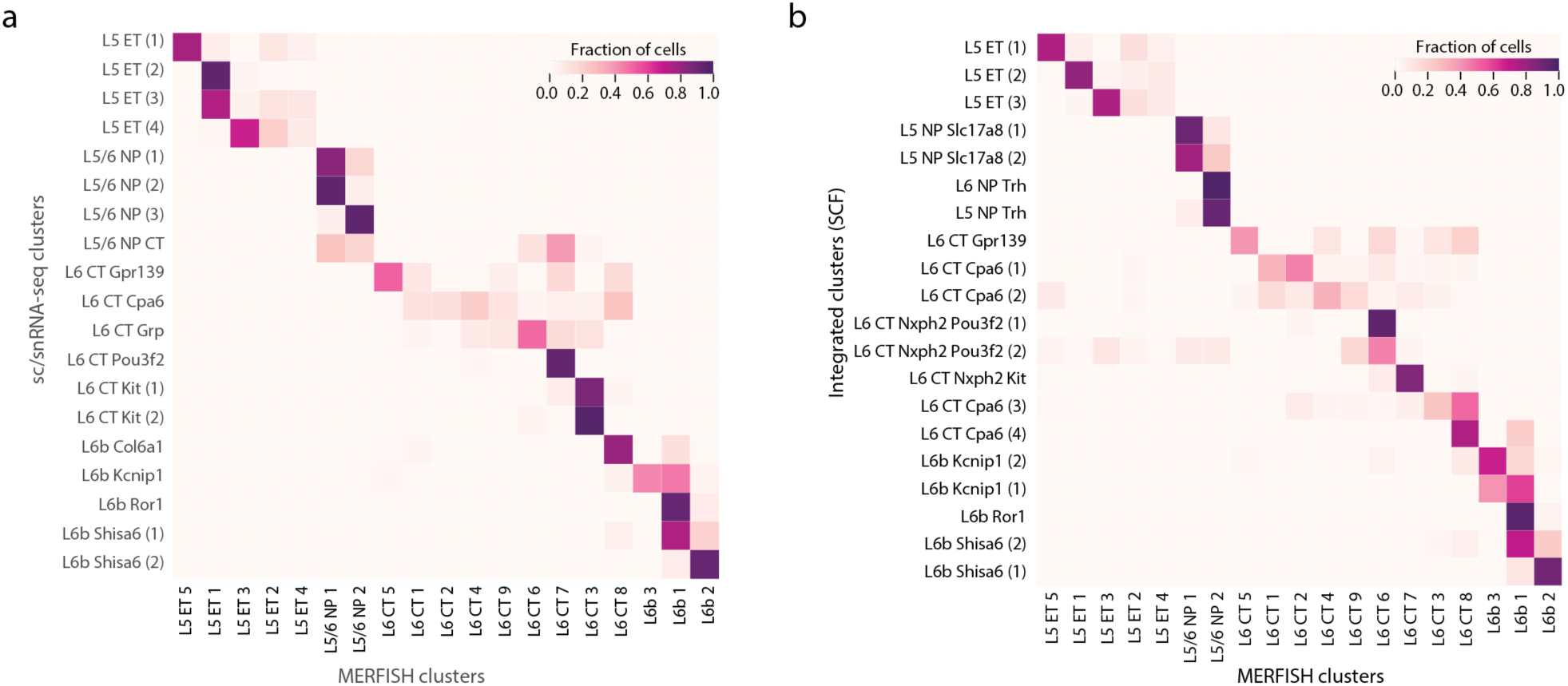
Correspondence between the ET, CT, NP and L6b clusters determined by MERFSH and those determined sc/snRNA-seq or by the integrated cluster analysis of sc/snRNA-seq, snATAC-seq and snmC-Seq. **a**, Correspondence between the ET, CT, NP and L6b clusters determined by MERFISH and by sc/snRNA-seq. **b**, Correspondence between the ET, CT, NP and L6b clusters determined by MERFISH and by integrated analysis of sc/snRNA-seq, snATAC-seq and snmC-Seq datasets using SingleCellFusion^38^. The classifier approach used to determine the correspondence is as described in Extended Data Figure 2.

**Extended Data Figure 6.**
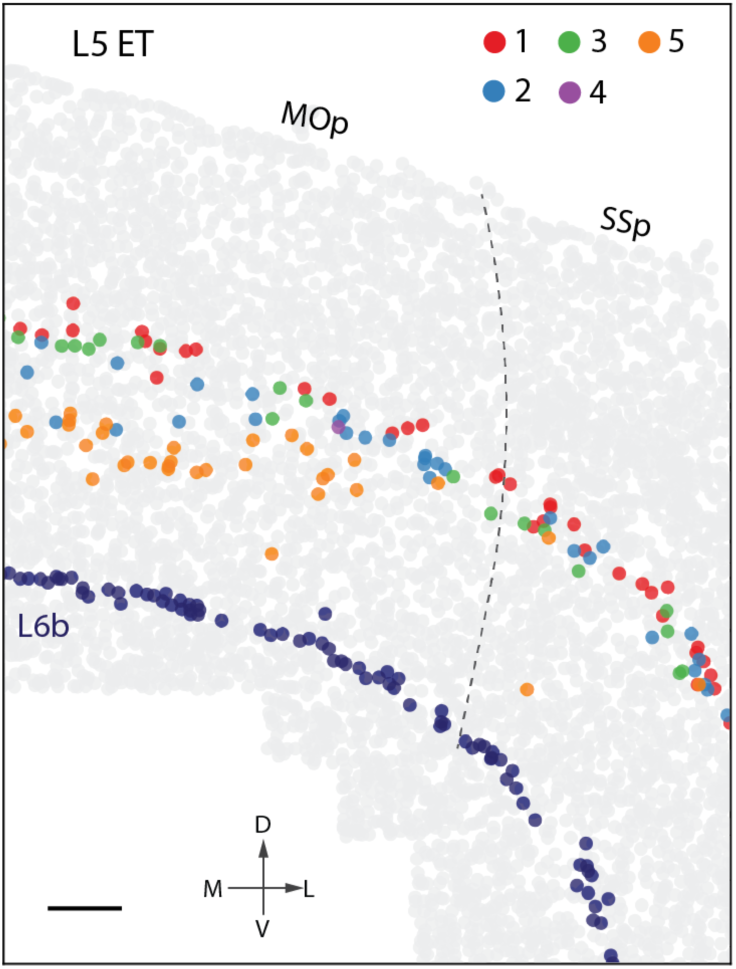
Different spatial distributions of ET cells in MOp and SSp. A coronal slice (∼Bregma +0.7) highlighting the L5 ET cells colored by cell clusters in the MOp and the neighboring SSp region. Dashed grey line marks the approximate border between MOp and SSp according to Allen CCF v3 (http://atlas.brain-map.org/). The L6b cells are shown in dark blue to mark the bottom boarder of the cortex.

**Extended Data Figure 7.**
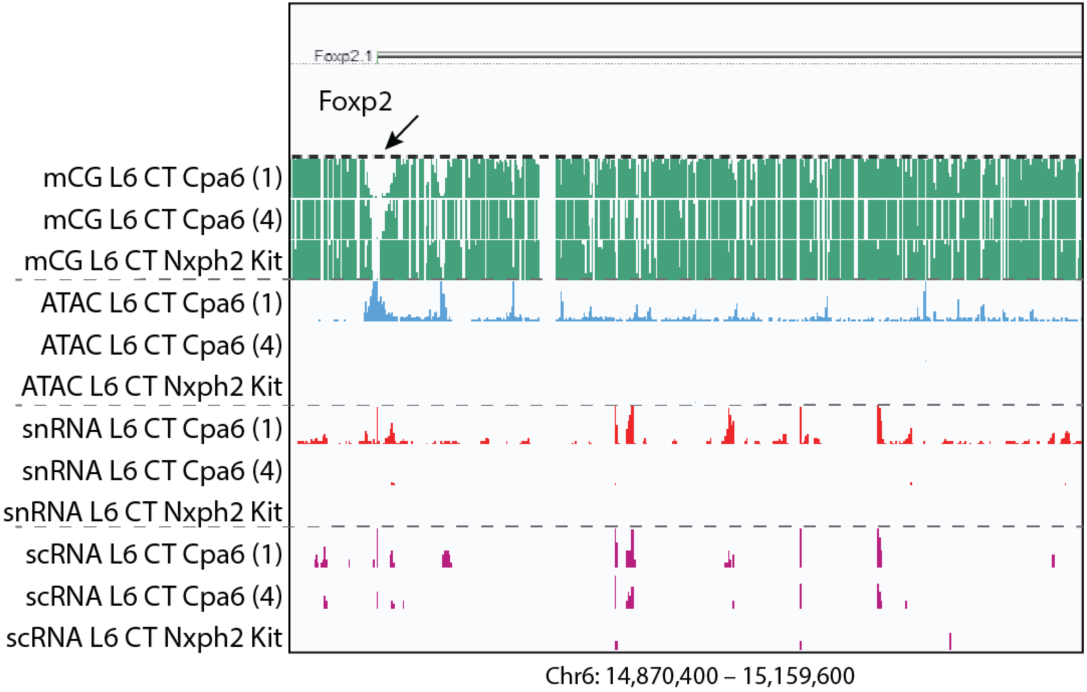
Epigenetic properties of representative marker genes for MERFISH clusters revealed by correspondence between clusters identified by MERFISH and integrated analysis. A marker gene for MERFISH cluster L6 CT 1, Foxp2, which also marks the corresponding integrated cluster L6 CT Cpa6 (1), shows enrichment in open chromatin reads and RNA counts, and moderate reduction in DNA methylation (indicated by an arrow) as compared to L6 CT Cpa6 (4) and Nxph2 Kit, which correspond MERFISH clusters L6 CT 8 and 7, respectively.

**Extended Data Figure 8.**
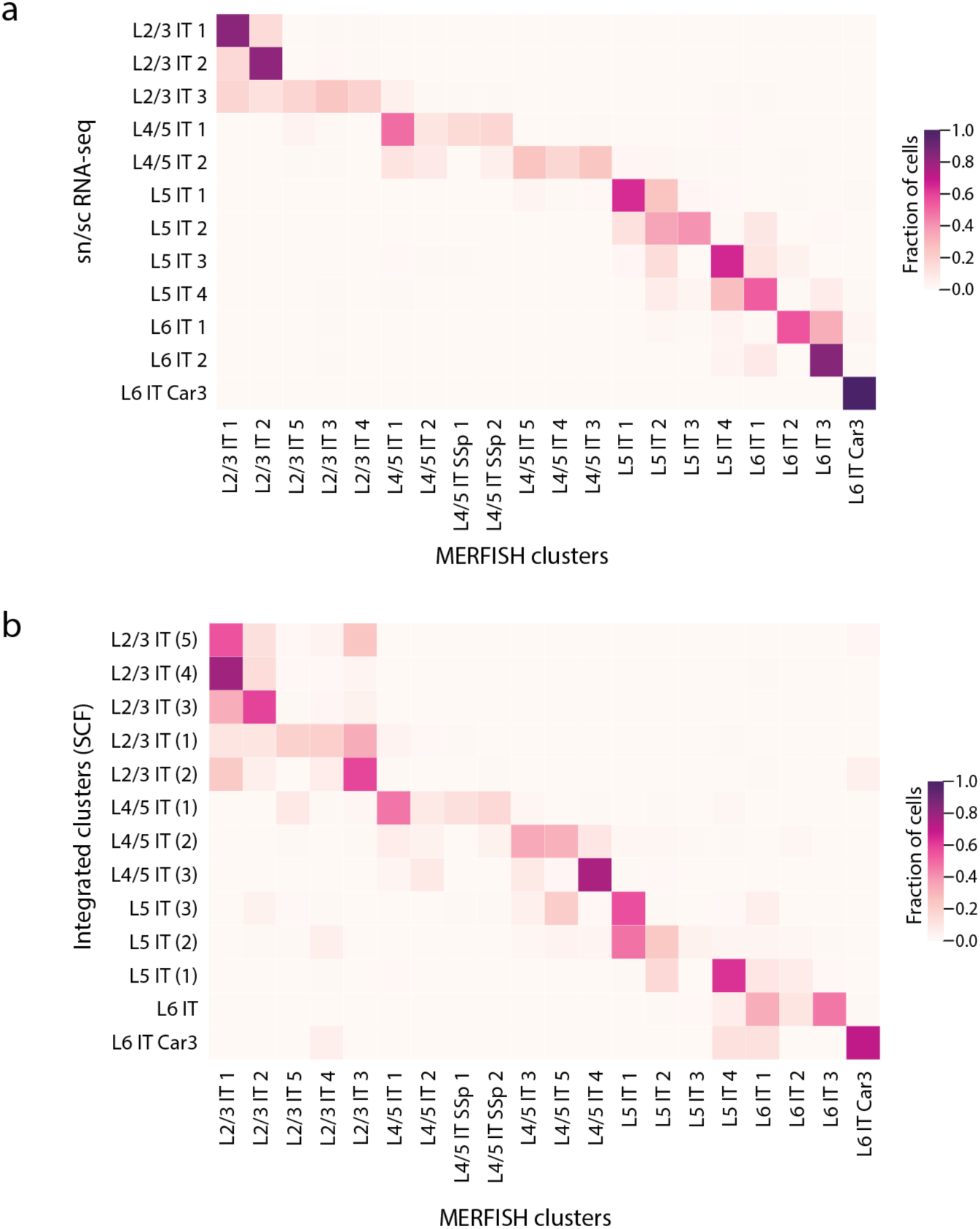
Correspondence between the MERFISH IT clusters and the clusters determined by sc/snRNA-seq analysis and those by integrated analysis. **a**, Correspondence between the IT clusters identified by MERFISH and those identified by the sc/snRNA-seq. **b**, Correspondence between the IT clusters identified by MERFISH and those identified by the integrated clustering analysis of sc/snRNA-seq, snATAC-seq and snmC-seq using SingleCellFusion^38^. The correspondence is determined using a classifier approach as described in Extended Data Figure 2.

**Extended Data Figure 9.**
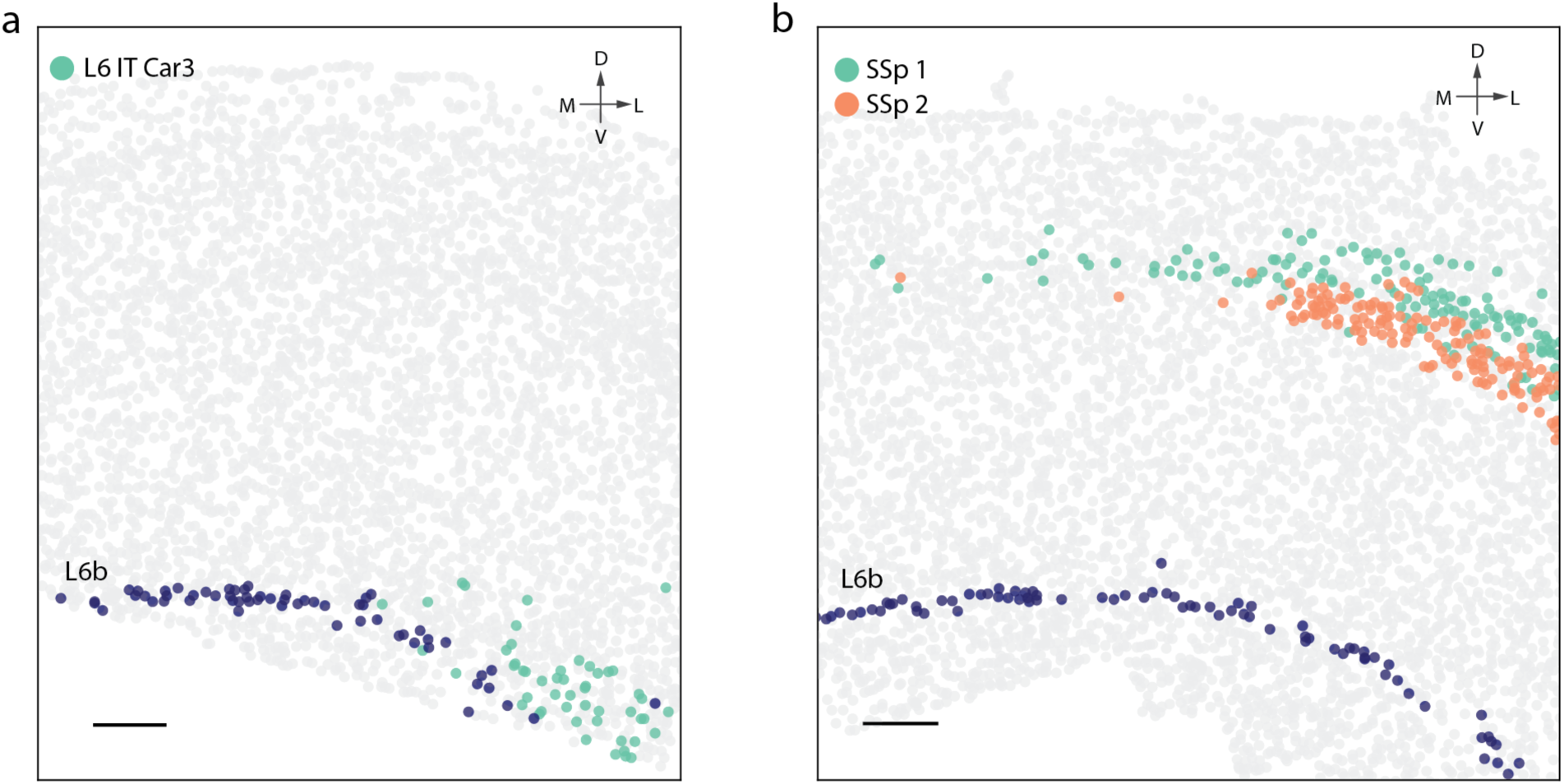
Spatial distributions of L6 IT Car3 and L4/5 IT SSp 1 and 2 clusters. **a**, A coronal slice (∼Bregma +1.2) highlighting the L6 IT Car3 cluster (green) shows that these cells are preferentially located at the lateral side of MOp in deep layer 6. **b**, A coronal slice (∼Bregma +0.7) highlighting the L4/5 IT SSp 1 (green) and L4/5 IT SSp 2 clusters (orange). The two SSp clusters form two dense layers in the SSp region and also extend into MOp. In both (a) and (b), the L6b cells are shown in dark blue to mark the bottom boarder of the cortex.

**Extended Data Figure 10.**
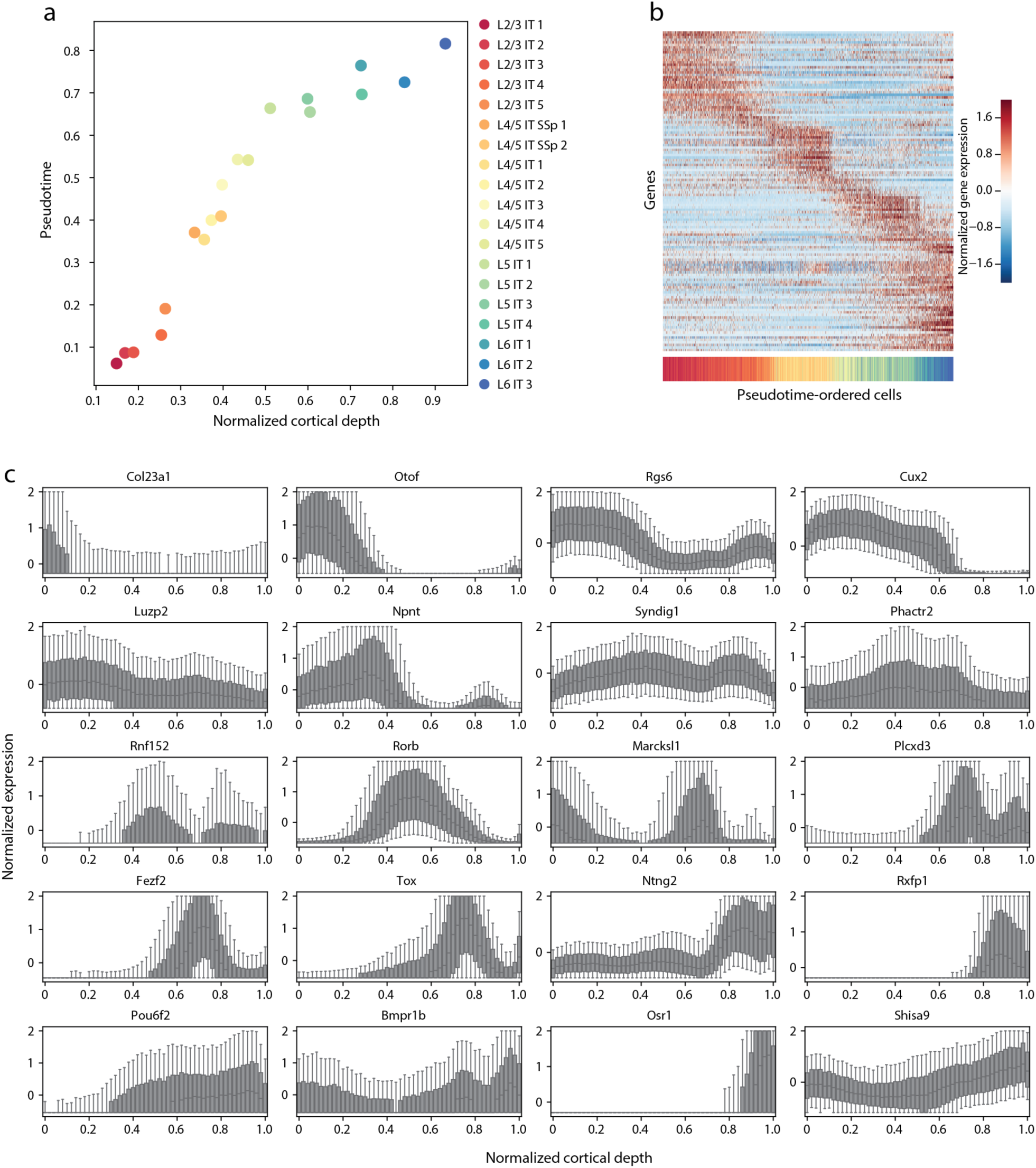
Correlation between gene expression and cortical depth position for IT cell clusters, and the dependence of expression levels of differentially expressed genes on pseudotime and cortical depth. **a**, Scatter plot of the mean pseudotime vs. mean normalized cortical depth for the IT clusters. **b**, Normalized expression of 128 genes that are differentially expressed across pseudotime. All IT cells were sorted in order of ascending pseudotime value, and the genes were sorted based on the pseudotime at which they exhibit maximal expression. The color bar beneath the heatmap indicates the cluster identity of the cell in that column of the heatmap, according to the color-cluster correspondence legend as shown in (a). The expression for each gene was z-score normalized across all IT cells, then the IT cells were grouped into 50 equal-sized bins according to their pseudotime and the mean normalized expression value of each bin is shown. **c**, Box plot of expression level in IT cells grouped into bins based on cortical depth, showing 20 representative genes that are differentially expressed across cortical depth. Expression was normalized and IT cells were grouped into 50 equal-sized bins according to their cortical depth. For each bin, the distribution of expression for the constituent cells are shown. The box represents the interquartile range, the median is indicated with a line, whiskers are 10th and 90th percentile, outliers are not shown.

## Online Methods

### Animals

Adult C57BL/6 male mice aged 57-63 days were used in this study. Animals were maintained on a 12 hour:12 hour light/dark cycle (2pm-2am dark period) with ad libitum access to food and water. Animal care and experiments were carried out in accordance with NIH guidelines and were approved by the Harvard University Institutional Animal Care and Use Committee (IACUC).

### Gene selection for MERFISH

In order to discriminate transcriptionally distinct cell populations with MERFISH, we designed a panel of 258 genes. Among the 258 genes, 62 were manually picked marker genes including established markers for inhibitory neurons and excitatory neurons, as well as different non-neuronal cell markers for mature and immature oligodendrocytes, astrocytes, microglia, macrophages, endothelial cells, pericytes, smooth muscle cells, and vascular leptomeningeal cells (VLMCs). To discriminate different neuronal types, we took two approaches to select genes. In the first approach, we selected a panel of most informative genes using mutual information analysis as reported previously^39^. Briefly, we used an information theory quantity known as mutual information^40^ to determine the relative amount of information each gene carries in defining the sc/snRNA-seq clusters. The amount of information that each gene carries is defined as the information gain due to knowledge of the expression state of the given gene (i.e. the mutual information between the gene and the cell classification). We used the scRNA 10x v2 A dataset generated by a companion study^38^ and determined highly variable genes using the Scanpy^65^ package. We binarized the expression profiles using a gene counts cutoff of zero to simplify the calculation of the mutual information. We selected the top 50 most informative genes based on mutual information analysis for excitatory neuronal clusters and inhibitory neuronal clusters, respectively, and due to overlap between the two groups, this approach generated a total of 91 top mutual information genes. In the second approach, we selected a panel of 168 genes based on differential expressed (DE) gene analysis using the scRNA-seq data (scRNA 10x v2 and scRNA SMART data) from the companion study^38^. We first found DE genes for each neuronal cluster pair (consisting a foreground cluster and a background cluster) in both directions. The criteria to define DE genes were: the genes have ≥2-fold change in expression between the foreground and background clusters and P-value < 0.05; they express in at least 40% cells in the foreground cluster, with more than 3-fold enrichment, in terms of the fraction of cells expressing the gene, relative to the background cluster. P-values were calculated using ANOVA test in limma^66^ on log transformed data. Top 50 genes that passed all the tests and ranked by P-values in each direction for every cluster pair were pooled together as candidates for scoring for the final marker set. To determine the final marker list, which we required to include at least two genes in both direction for all pairs of clusters, we used a greedy algorithm to find minimal number of genes that satisfy the requirement. Starting from the 62 manually picked marker list as described above, the algorithm checks which pairs already have sufficient number of DE genes, and work on the remaining pairs of clusters until each pair of clusters has 2 DE genes included in both directions. This approach generated a total of 168 genes.

We then combined the marker lists generated by these three different approaches, which partially overlap with each other, resulting in a panel of 258 genes total. We then screened this gene list to identify genes that are relatively short or have relatively high expression level, which were potentially challenging for highly multiplexed FISH imaging experiments as described previously^36^. We found 16 genes that can accommodate fewer than 48 hybridization probes with target sequences that are 30-nucleotide (nt) long, or are expressed at an average of 200 counts in any cell cluster as determined from the scRNA SMART data^38^. These 16 genes were imaged in a set of eight sequential, two-color FISH imaging rounds, following the MERFISH run that imaged the remaining 242 genes.

### Design and construction of the MERFISH encoding probes

MERFISH encoding probes for the 242 genes were designed as previously described^36^. We first assigned to each of the 242 genes a unique binary barcode drawn from a 22-bit, Hamming-Distance-4, Hamming-Weight-4 encoding scheme. We included 10 extra barcodes as “blank” barcodes, which were not assigned to any genes, to provide a measure of the false-positive rate in MERFISH as described previously^36^.

We identified all possible 30-mer targeting regions within each desired gene transcript as previously described^67^. Each MERFISH encoding probe contains a 30-mer targeting region that is complementary to the RNA of interest, as well as two 20-mer readout sequences that encode the specific barcode assigned to each gene. From the set of all possible 30-mer probes for each gene, we selected 92, 30-mer probes at random. For the transcripts that were not long enough and had fewer than 92 probes, we allowed these 30-mers to overlap by as much as 20 nt to increase the number of probes – because a given cellular RNA is typically bound by less than one third of the 92 encoding probes^68^, the encoding probes with overlapping targeting regions do not substantially interfere with each other but partially compensate for reduced binding due to local inaccessible regions on the target RNA or loss of probe during synthesis. We then assigned two readout sequences to each of the encoding probes associated with each gene. For the 22-bit encoding scheme, a total of 22 readout sequences were used, each associated with one bit, and the collection of encoding probes for each gene together contain 4 of the 22 readout sequences that corresponded to the 4 bit that reads “1” in the barcode assigned to that gene.

Encoding probes for the 16 genes imaged in sequential two-color FISH rounds were produced in the same fashion, except that one single unique readout sequence was concatenated to each of these probes. The readout sequences used here were different from the 22 readout sequences used for MERFISH run.

In addition, we concatenated to each encoding probe sequence two PCR primers, the first comprising the T7 promoter, and the second being a random 20-mer designed to have no region of homology greater than 15 nt with any of the encoding probe target sequences designed above, as we previously described^67^.

With the template encoding probe sequences we designed above, we constructed the MERFISH probe set as previously described^36^. The template DNA were synthesized as a complex oligo pool (Twist Biosciences). This pool contained both the encoding probes to the 242 genes measured in MERFISH run and the 16 genes measured in sequential two-color FISH rounds, but different primer sequences, which allowed us to amplify these two groups separately via PCR followed by same synthesis and purification procedures. The two groups were then mixed during tissue staining.

### Design and construction of MERFISH readout probes

For the 258 gene panel used in this study, 38 readout probes were designed, each complementary to one of the 38 readout sequences. 22 of the 38 readout probes correspond to the 22 bits barcodes used for MERFISH imaging, and the remaining 16 readout probes each corresponds to one gene that was imaged in the sequential two-color FISH rounds. Each readout probes were conjugated to one of the two dye molecules (Alexa750, Cy5) via a disulfide linkage, as described previously^67^. These readout probes were synthesized and purified by Bio-synthesis, Inc., resuspended immediately in Tris-EDTA (TE) buffer, pH 8 (Thermo Fisher) to a concentration of 100 µM and stored at -20 °C.

### Tissue preparation for MERFISH

Mice aged 57-63 days were euthanized with CO_2_, and their brain were quickly harvested and cut into hemispheres and each hemisphere was frozen immediately on dry ice in optimal cutting temperature compound (Tissue-Tek O.C.T.; VWR, 25608-930), and stored at -80°C until cutting. Frozen brain hemispheres were sectioned at -18°C on a cryostat (Leica CM3050 S). Slices were removed and discarded until the MOp region was reached. A continuous set of 300, 10-µm-thick slices were cut from anterior to posterior, and approximately every 10^th^ slice was placed onto coverslips for imaging. Each coverslip contained 5-6 slices. In total 32 slices were collected for each animal. The coverslips were prepared as described previously^35,36^.

Tissue slices were fixed by treating with 4% PFA in 1×PBS for 15 minutes and were washed three times with 1×PBS and stored in 70% ethanol at 4 °C for at least 18 hours to permeabilize cell membranes. The tissue slices from the same animal were cut at the same time and distributed to six coverslips, which were store in 70% ethanol at 4 °C for no longer than 2 weeks until all the coverslips were imaged. We observed no degradation in sample quality over this time.

The tissue slices were stained with the MERFISH probe set as described previously^36^. Briefly, the samples were removed from the 70% ethanol and washed with 2× saline sodium citrate (2×SSC) for three times. Then we equilibrated the samples with encoding-probe wash buffer (30% formamide in 2×SSC) for five minutes at room temperature. The wash buffer was then aspirated from a coverslip, and the coverslip was inverted onto a 50 µL droplet of encoding-probe mixture on a parafilm coated petri dish. The encoding-probe mixture comprised ∼1 nM of each encoding probe for the MERFISH run, ∼5 nM of each encoding probe for the sequential two-color FISH rounds, and 1 µM of a polyA-anchor probe (IDT) in 2×SSC with 30% v/v formamide, 0.1% wt/v yeast tRNA (Life Technologies, 15401-011) and 10% v/v dextran sulfate (Sigma, D8906). We then incubated the sample at 37 °C for 36∼48 hours. The polyA-anchor probe containing a mixture ofDNAandLNAnucleotides (/5Acryd/TTGAGTGGATGGAGTGTAATT+TT+TT+TT+TT+TT+TT+TT+TT+TT+T, were T+ is locked nucleic acid, and /5Acryd/ is 5’ acrydite modification) hybridized to the polyA sequence on the polyadenylated mRNAs and allowed these RNAs to be anchored to a polyacrylamide gel as described below. After hybridization, the samples were washed in encoding-probe wash buffer for 30 minutes at 47 °C for a total of two times to remove excess encoding probes and polyA-anchor probes. All tissue samples were cleared to remove fluorescence background as we previously described^35,36^. Briefly, the samples were embedded in a thin polyacrylamide gel and were then treated with a digestion buffer of 2% v/v sodium dodecyl sulfate (SDS; ThermoFisher, AM9823), 0.5% v/v Triton X-100 (Sigma, X100), and 1% v/v proteinase K (New England Biolabs, P8107S) in 2×SSC for 36-48 hours at 37 °C. After digestion, the coverslips were washed in 2×SSC for 30 minutes for a total of four washes and then stored at 4°C in 2×SSC supplemented with 1:100 Murine RNase inhibitor (New England Biolabs, M0314S) prior to imaging.

### MERFISH imaging

We used a home-built imaging platform in this study as previously described^34^. To prepare the sample for imaging, we first stained it with a readout hybridization mixture containing the readout probes associated with the first round of imaging in the MERFISH run, as well as a probe complementary to the polyA-anchor probe and conjugated via a disulfide bond to the dye Alexa488 at a concentration of 3 nM. The readout hybridization mixture was comprised of the readout-probe-wash buffer comprised of 2×SSC, 10% v/v ethylene carbonate (Sigma, E26258), and 0.1 % v/v Triton X-100, supplementing with 3 nM each of the appropriate readout probes. The sample was incubated in this mixture for 15 minutes at room temperature, and then washed in the readout-probe wash buffer supplemented with 1 µg/mL DAPI for 10 minutes to stain nuclei within the sample. The sample was then washed briefly in 2×SSC and imaged. Briefly, the sample was loaded into a commercial flow chamber (Bioptechs, FCS2) with a 0.75-mm-thick flow gasket (Bioptechs, 1907-100; DIE# F18524). Imaging buffer comprising 5 mM 3,4-dihydroxybenzoic acid (Sigma, P5630), 2 mM trolox (Sigma, 238813), 50 µM trolox quinone, 1:500 recombinant protocatechuate 3,4-dioxygenase (rPCO; OYC Americas), 1:500 Murine RNase inhibitor, and 5 mM NaOH (to adjust pH to 7.0) in 2×SSC was introduced into the chamber and the sample was imaged with a low magnification objective (Nikon, CFI Plan Apo Lambda 10x) with 405-nm illumination to produce a low-resolution mosaic of all slices in the DAPI channel. We then used this mosaic image to locate the MOp region in each slice and generated a grid of field-of-view (FOV) positions to cover the MOp region to be imaged. We then switched to a high magnification, high-numerical aperture objective (Nikon, CFI Plan Apo Lambda 60x) and imaged each of the FOV positions generated above. In the first round of imaging, we collected images in the 750-nm, 650-nm, 560-nm, 488-nm, and 405-nm channels to image the first two readout probes (conjugated to Alexa750 and Cy5, respectively), the orange fiducial beads, the total polyA-mRNA stained by the polyA-anchor probe (Alexa488), and the nucleus stained by DAPI (405-nm channel). The latter two channels were used for cell segmentation as described below. We took a single image for the fiducial beads on the surface of the coverslip using the 560-nm illumination channel for each imaging round as a spatial reference to correct for slight misalignments in the stage position over the imaging rounds. To image the entire volume of each 10-µm-thick slice, we collected seven 1.5-µm-thick z-stacks for other four channels (two readout probes, polyA probe and DAPI) in each FOV.

After the first round of imaging, the dyes were removed by flowing 2.5 mL of cleavage buffer comprising 2× SSC and 50 mM of Tris (2-carboxyethyl) phosphine (TCEP; Sigma, 646547) with 15 min incubation in the flow chamber, in order to cleave the disulfide bond linking the dyes to the readout probes. The sample was then washed by flowing 1.5 mL 2× SSC.

To perform subsequent rounds of imaging, we flowed 3.5 mL of the readout probe mixture containing the appropriate readout probes across the chamber and incubated the sample in this mixture for a total of 15 minutes for each round. Then the sample was then washed by 1.5 mL of readout-probe wash buffer and then 1.5 mL of imaging buffer was introduced into the chamber. For each round, we took images for all FOV locations in the 750-nm, 650-nm, and 560-nm channels for the 2 readout probes and fiducial beads. Two readout probes were imaged in each round, one labeled with Alexa750and the other with Cy5, and a readout probe mixture containing 3 nM of appropriate readout probes was used for each round. We repeated the hybridization, wash, imaging and cleave for all rounds to complete the 22-bit MERFISH imaging and the 8 rounds of sequential two-color FISH. All buffers and readout probe mixtures were loaded with a home-built, automated fluidics system composed of three, 12-port valves (IDEX, EZ1213-820-4) and a peristaltic pump (Gilson, MP3), configured as previously described^7^.

### MERFISH image analysis and cell segmentation

All MERFISH image analysis was performed using MERlin^69^, a Python-based MERFISH analysis pipeline, using algorithms similar to what we have described previously^34,36^. First, we aligned the images taken during each imaging round based on the fiducial bead images, accounting for X-Y drift in the stage position relative to the first round of imaging. For the MERFISH images, we then high-pass filtered the image stacks for each FOV to remove background, deconvolved them using 20 rounds of Lucy-Richardson deconvolution to tighten RNA spots, and low-pass filtered them to account for small movements in the apparent centroid of RNAs between imaging rounds. Individual RNA molecules were identified by our previously published pixel-based decoding algorithm^67^. After assigning barcodes to each pixel independently, we aggregated adjacent pixels that were assigned with the same barcodes into putative RNA molecules, and then filtered the list of putative RNA molecules to enrich for correctly identified transcripts as described previously^34^ for a gross barcode misidentification rate at 5%. We further removed putative RNAs that contained only a single pixel as they are prone to be background of spurious barcodes generated by random fluorescent fluctuations and had a much higher misidentification rate compared to those contained 2 and more pixels.

We identified cell segmentation boundaries in each FOV using a seeded watershed approach as described before^36^. The DAPI images were used as seeds and the polyA signals were used to identify segmentation boundaries. Finally, we assigned individual RNA molecules identified in the MERFISH run to individual cells based on whether or not they fell within the segmented boundaries of the cells. For the sequential two-color FISH rounds, we quantified the signal from these imaging rounds by summing the fluorescence intensity of all pixels that fell within the segmentation boundaries of the cells associated with the central z-plane and normalized the signal by the areas of the cells in this z-plane. Then the normalized signals of the 16 genes from the sequential two-color FISH rounds were merged with the RNA counts matrix from the 242 genes measured in MERFISH run and used for cell clustering analysis.

### Cell clustering analysis of MERFISH data

With the cell-by-gene matrix obtained as described above (each row representing a cell and each column representing a gene, and each element representing the expression level a specific gene in a specific cell), we first preprocessed the matrix by several steps. (1) The segmentation approach we used generated a small fraction of putative “cells” with very small total volumes due to spurious segmentation artifacts, as well as some cells that overlapped in the 3-D dimension and were not properly separated. We hence removed the segmented “cells” that had a volume that was either less than 100 µm^3^ or larger than 3 times of the median volume of all cells, which was about 1000 µm^3^. (2) A fraction of cells did not have the whole soma body included in a 10-µm-thick tissue slice and were thus not imaged completely. To remove the differences in RNA counts due to the incompleteness of soma bodies, we normalized the RNA counts per cell by the imaged volume of each cell. (3) We observed a modest batch effect between MERFISH experiments accounting for ∼30% variation of the mean total number of RNAs per cell. We normalized the mean total RNA counts per cell to a same mean value (250 in this case) for each experiment to remove the influence of these batch effects. (4) Since the 16 genes that were imaged in the sequential FISH rounds contained many non-overlapping markers and no cells should express a majority of these 16 genes, we considered the segmented “cells” that had a normalized fluorescence signal that higher than the 90% quantile in 12 out of the total 16 sequential FISH channels as caused by spurious fluorescence background and removed these “cells”. (5) Since the fluorescence background in the 650-nm and 750-nm channels were different, we subtracted the background for each cell by taking the minimum of the signal for each cell across all sequential FISH rounds for 650-nm and 750-nm channels separately. (6) We removed the cells that had total RNA counts lower than 2% quantile or higher than 98% quantile. (7) We removed potential doublets using Scrublet^70^. Briefly, principal component analysis (PCA) was used to train a k-nearest neighbor (kNN) classifier to predict a doublet score for each cell. Since we recorded the DAPI stained nucleus image of each cell, we were able to visually inspect a random subset of potential doublets picked by Scrublet and fine-tuned the doublet score threshold to remove connected cells more accurately. Finally, the cells with doublet score higher than 0.18 were removed as doublets, which accounted for ∼12% of the total cell number. (8) We also found that 4 out of the 16 genes imaged in the sequential two-color runs, namely *Cd52, Rprml, Mup5* and Igfbp6, were not detected well in all experiments and failed to mark any subset of cells as revealed in following clustering analysis. These 4 genes were removed for subsequent analysis.

After the above preprocessing steps, we normalized the total RNA counts for each cell to the median total RNA counts of all cells and log transformed the cell-by-gene matrix. We then normalized their expression profiles by computing the z-score for each gene. We performed dimensionality reduction of the matrix using PCA, and used the first 35 principal components (PCs). To determine the number of PCs to keep, we calculated the largest eigenvalue from a randomly shuffled values in each column of the cell-by-gene matrix, and then kept all the PCs that had an eigenvalue higher than the mean of the largest eigenvalues across 20 iterations of random shuffling. We then performed graph-based Louvain community detection^42^ in the 35 PC space using Scanpy^65^ for a range of nearest neighborhood size k values with a bootstrap analysis to both identify stable clusters and select the optimal k value (k = 10) as described previously^36^. We further identified six small clusters that expressed mixtures of markers for multiple distinct cell classes, e.g. *Slc17a7* which marks excitatory neurons and *Sox10* which marks the oligodendrocytes, and that did not correspond to any of the major subclasses defined by the sc/snRNA-seq data^38^ (based on classifier analysis which will be described below), as potential doublets, which were excluded from subsequent analysis.

From the first round of clustering, we identified 16 excitatory neuronal clusters, 8 inhibitory neuronal clusters and 14 other clusters. To further refine our detection of transcriptionally distinct populations, we separated all the cells into five groups: intra-telencephalic (IT) projecting neurons (marked by excitatory neuronal marker *Slc17a7* and pan IT marker *Slc30a3*), non-intra-telencephalic (non-IT) neurons (marked by excitatory neuronal marker *Slc17a7* but not *Slc30a3*), caudal ganglionic eminence (CGE) derived inhibitory neurons (marked by *Gad1, Gad2*, and *Lamp5/Sncg/Vip*), medial ganglionic eminence (MGE) derived inhibitory neurons (marked by *Gad1, Gad2*, and *Sst/Pvalb*), and other cells. We then repeated the procedure of dimensionality reduction and clustering, as described above, for these five cell groups separately. In addition, we sampled a range of resolution parameter r (r=1, 2, 3), a parameter value defined in Scanpy^65^ that controls the coarseness of the clustering, to search for the best granularity that represent the diversity of the transcriptomic profiles. We kept k=40 and r=2 for IT and non-IT excitatory neurons, k=15 and r=2 for CGE and MGE derived inhibitory neurons, and k=20, r=1 for the non-neuronal cells.

After the second round of clustering, we further removed a small fraction of cells as potential doublets as described above. We also found four unique clusters that didn’t correspond to any subclass in the MOp region defined by the sc/snRNA-seq data^38^ (using the classifier approach described below). We located the cells that belonged to these clusters and found that two clusters were in striatum and were probably striatum neurons, and the other two clusters were likely ependymal cells locating in the lateral ventricle. We removed these clusters from subsequent analysis.

For presentation, Uniform Manifold Approximation and Projection (UMAP)^52^ was used to embed the cells in two dimensions using the same PCs that were used for clustering.

### A neural-net classifier approach to determine correspondence between clusters identified by MERFISH and sequencing-based measurements

Correspondence between cell clusters identified by MERFISH and by sc/snRNA-seq were assessed by running a neural-net classifier^71^ which was trained on the z-scored single-cell expression profiles measured by MERFISH. The snRNA-seq 10x v3 B data in the companion paper^38^ was used for comparison because it is the largest dataset among the seven single cell and single nucleus RNA-seq datasets included in this companion study and contained the most non-neuronal cells, while all other six datasets were collected by fluorescence-activated cell sorting (FACS) to enrich neurons. The snRNA 10x v3 B data was z-scored, and then the subset of genes measured in the MERFISH data were used together with the trained model to predict a MERFISH cluster label for each cell in the snRNA-seq dataset. From this, each snRNA-seq cell had both a predicted MERFISH cluster label and a cluster label determined from the consensus clustering results for the seven sc/snRNA-seq datasets^38^. Cells were grouped based on their consensus sc/snRNA-seq cluster identity, and then the fraction of cells from a given consensus sc/snRNA-seq cluster that were predicted to have each MERFISH cluster was then determined. Such a classifier can also be trained on the snRNA-seq dataset and used for predicting a sc/snRNA-seq cluster label for each cell in the MERFISH dataset. We obtained similar results by running the classifier both ways, but only presented the results from the classifier trained by the MERFISH data. The same classifier approach was also used to produce Extended Data Figure 2a but in this case the subclass labels defined by MERFISH and sc/snRNA-seq data for each cell was used instead of cluster labels. Likewise, the same classifier approach was used to produce Extended Data Figures 2a, 5b and 8b, but in these cases, the cluster labels defined by the integrated analysis of the seven sc/snRNA-seq datasets, a snATAC-seq dataset and a snmC-seq dataset were used instead of the cluster labels derived from the sc/snRNA-seq datasets alone.

### Analysis of gene expression and spatial continuity among IT cells

To visualize the degree of similarity and continuity in the gene expression profiles of the IT neuronal clusters, we employed a recently developed graph abstraction technique called PAGA^53^ to gain a quantitative understanding of how extensively different IT cell clusters occupied overlapping gene expression space. To this end, we first took the 19 IT neuronal clusters and normalized their expression profiles by computing the z-score for each gene. Cells from the L6 IT Car3 were not included in this analysis as it formed a cluster that was well-separated in gene expression from the other IT cell clusters. PCA was used to reduce dimensionality of the normalized expression data to the first 19 PCs. In selecting the number of PCs to include, we performed the same randomization procedure used when setting a PC threshold during clustering as described in the “Cell clustering analysis of MERFISH data” above. We then constructed a kNN graph based on the PCs, identifying the 12 nearest neighbors of each cell. Using the kNN graph and the cluster label of each cell, we used Scanpy^65^ to calculate the frequency that edges from cells with a given cluster label were connected to cells from a different cluster label and then normalize this frequency to that expected by chance. The resulting values represent the connectivity between the clusters in the kNN graph, and are visualized in a graph wherein each cluster is a node and the edges between nodes indicate the connectivity between those clusters.

Next, we constructed an ordering of the IT cells based on their expression profile, yielding a “pseudotime” value for each cell. This calculation is most often performed to order cells within a dynamic system, in which case the ordering reflects the “time” relative to some reference cell. This calculation performed on the IT cells is not intended to represent the trajectory from L2/3 to L6 as part of a dynamic process, but rather to obtain an expression-derived measure of where along the trajectory each cell falls. To calculate the pseudotime of the IT cells, we used Scanpy to construct a diffusion map based on the above-described kNN graph, assigned a neuron from the L2/3 IT 1 cluster as the root cell of the trajectory, and then computed the diffusion-based pseudotime^72^. The resulting value assigned to each cell reflects how far from the root cell its expression profile places it, and since each cell falls along a single trajectory with the L2/3 IT root cell at one end, this value orders the cells relative to one another along this path.

To identify genes that vary as a function of IT cell pseudotime, the expression profile of the IT cells was normalized by computing the z-score for each gene. The IT cells were split evenly into 50 bins based on their pseudotime rank-order, and the mean normalized expression was calculated for each gene across all the bins. Any gene for which the difference in mean normalized expression between any two bins exceeded 0.5 was selected as a differentially expressed gene. For plotting these genes in a heatmap, the genes were ordered according to the pseudotime at which they exhibit their maximum expression and the cells were ordered based on their pseudotime. A rolling average was calculated for each gene across the 50 bins, using a window size of 10 bins. Genes were then sorted according to the window at which the maximum value occurred. The same procedure was used to find and sort genes that vary as a function of the cortical depth values of the IT cells, using normalized cortical depth in place of pseudotime.

### Stereotaxic injection of retrograde tracers

To retrogradely label TEa-, SSp-, and MOs-projecting MOp neurons, each region was injected in the same animal in the right hemisphere with 100 nL of fluorescently conjugated Cholera Toxin subunit b (CTb-AlexaFluor488, CTb-AlexaFluor555, or CTb-AlexaFluor647, respectively; 0.5%; ThermoFisher, Cat# C22841, C22843, and C34778) using the following coordinates relative to bregma: TEa (AP -1.7 mm, ML +4.5 mm, DV +2.5 mm below cortical surface), SSp (AP -0.5 mm, ML +2.4 mm, DV +0.5 mm below cortical surface), and MOs (AP +2.4 mm, ML +1.0 mm, DV +0.4 mm below cortical surface). Injection procedures were performed in adult male and female C57BL/6J mice (Jackson Laboratories) aged 2-4 months. Briefly, mice were anesthetized initially in an induction chamber containing 5% isoflurane mixed with oxygen and then transferred to a stereotaxic frame equipped with a heating pad. Anesthesia was maintained throughout the procedure using continuous delivery of 2% isoflurane through a nose cone at a rate of 1.5 L/min. The scalp was shaved, and a small incision was made along the midline to expose the skull. After leveling the head relative to the stereotaxic frame, the specified injection coordinates were used to mark the locations on the skull directly above each target area and a small hole (0.5 mm diameter) was drilled for each. CTb was delivered through pulled glass micropipettes (inner diameter of tip: ∼20 µm) using pressure injection via a micropump (World Precision Instruments, Sarasota, FL). After completing the last injection, the scalp was sutured closed and animals were administered ketofen (5 mg/kg) to minimize inflammation and discomfort. Animals were recovered from anesthesia on a heating pad and then returned to their home cage. Mice were euthanized 7 days following injection to allow time for tracer transport and fresh brain tissue was immediately extracted, embedded in Tissue-Tek O.C.T. Compound (Sakura, Cat# 4583), and frozen at -80°C for later cryostat sectioning.

### Imaging for CTb-injected tissue

Frozen CTb-injected mouse brain was sectioned the same as described in the “Tissue preparation for MERFISH” section. Continuous set of 10-µm-thick slices in the MOp upper limb region (Bregma +0.7 to +0.1) were sectioned with approximately every 2^th^ slice kept and placed onto coverslips for imaging. We used a much higher sampling frequency for CTb-injected samples due to a higher failure rate of this experiment caused by removing the coverslip from the flow chamber after CTb imaging. Tissue slices were immediately fixed by treating with 4% PFA in 1×PBS for 15 minutes, washed three times with 1×PBS, stained with DAPI and proceed for imaging. As described in the “MERFISH imaging” section, we used the same imaging buffer and the sample was first imaged with a low magnification objective (Nikon, CFI Plan Apo Lambda 10x) for DAPI in 405-nm channel to produce a low-resolution mosaic of all slices. Next, in order to align each cell in the tissue with the same tissue slice that will be imaged with the MERFISH probe set later, we picked 10 cells in each coronal slice and recorded the location of the right-side edge for each cell. We then used the mosaic image, created as described above, to locate the MOp region in each slice and generated a grid of FOV positions to cover the MOp region to be imaged. We then switched to the high magnification objective (Nikon, CFI Plan Apo Lambda 60x) and collected images in the 650-nm channel for CTb-AlexaFluor647, 560-nm channel for CTb-AlexaFluor555, 488-nm channel for CTb-AlexaFluor488, and 405-nm channel for DAPI. We took a single image for each of these channels at the central z-plane.

After the CTb signals were imaged, the sample was removed from the imaging chamber and washed three times by 2xSSC and then permeabilized by 70% ethanol at 4 °C for at least 18 hours to permeabilize cell membranes. The tissue slices were then stained with the same MERFISH probe set as described in the “Tissue preparation for MERFISH” section, followed by normal MERFISH preparation procedures and imaging as described in the “Tissue preparation for MERFISH” and “MERFISH imaging” sections. During MERFISH imaging, we first imaged DAPI again with a low magnification objective, and then located the same 10 cells in each coronal slice we selected before during CTb imaging, and recorded the new location of the right-side edge for each cell. Using the old and new locations of the 10 cells for each slice, we determined the rotation and translation of the tissue comparing to the original images to align the CTb and MERFISH images. Then the normal MERFISH imaging was proceeded and the MERFISH images were decoded and segmented the same as described in the “MERFISH image analysis and cell segmentation” section. We assigned each cell a projection identity by thresholding the normalized CTb dye intensity for each CTb channel and labeled each cell “on” or “off” for each channel. The CTb labeling of the cells were mostly binary (“on” or “off”) but still the labeling level varied between cells, therefore threshold was tuned by manually examining a random subset of the images and was set to a fairly stringent level such that weakly labeled cells were labeled “off”. The cell type identities of the CTb-injected samples were determined by training the MERFISH dataset with the MERFISH cell cluster identities without CTb injections using the classifier as described in the “A neural-net classifier approach to determine correspondence between clusters identified by MERFISH and sequencing-based measurements” section and predicting on the CTb-injected samples. Each cell in the CTb-injected samples was hence assigned with both a cell type identity and a projecting target identity.

### Other data resources used in this work

The datasets of different single-cell transcriptomic and epigenomic modalities (scRNA SMART, scRNA 10x v3 A, scRNA 10x v2 A, snRNA SMART, snRNA 10x v3 B, snRNA 10x v3 A, snRNA 10x v2 A, snATAC-seq, snmC-seq) were generated by a concurrent study in the BICCN consortium^37^ and reported in a companion paper^38^. These data are available at the Neuroscience Multi-omics Archive (nemoarchive.org).

